# Unified Multi-caller Ensemble (UME) generates an unbiased maize haplotype map for variable coverage whole genome data

**DOI:** 10.1101/2023.12.07.570552

**Authors:** Miguel Vallebueno-Estrada, Kelly Swarts

## Abstract

We present a novel diversity-focused haplotype map (HapMap) that characterizes over 64.5 million maize (*Zea mays* ssp*. mays*) single nucleotide polymorphisms (SNPs) genotyped across 818 individuals from diverse backgrounds. This HapMap aims to balance the variation obtained from domesticated landraces and inbred lines, outgroup *Zea spp.* and more distant *Tripsacum spp.* in order to minimize ascertainment bias for diversity studies. Included individuals derive from public data from various experimental setups and coverages, which is challenging for standard SNP callers to accommodate. We provide evidence of coverage biases associated with standard callers that influence resulting variation and introduce a novel approach called Unified Multi-Caller Ensemble (UME), which enhances variant calling accuracy in low-coverage and mixed-coverage genomic datasets. UME corrects for coverage bias resulting from inter-sample coverage heterogeneity by leveraging evidence from variant callers with orthogonal strategies, re-calibrating the error probabilities across callers to minimize the impact of error biases inherent to a given caller. It outperforms individual strategies and excels in *de novo* variant calling, taking advantage of instances of higher depth reads, even in low coverage individuals, while preserving biologically informative variant relationships across coverage levels. An important feature of UME is the independence from population allele frequencies in the discovery panel, thus avoiding ascertainment bias resulting from unbalanced input genetic diversity. Discovered variants are unbiased because no population filtering is used, and the full diversity of SNPs is retained in the final variant call set to maximize the utility of the dataset for production calling newly sequenced samples. We present a strategy for filtering the recalibrated error profiles that relies on maximizing demographic signals to retain genetic relationships within the population while reducing sequencing error. After the variant discovery phase, we employ the UME production stage, which enriches genotype calling across all coverage levels, benefiting low-coverage samples. Error introduced in this process is removed through subsequent filtering. Using this approach, we generated a coverage bias-controlled maize HapMap database, providing a comprehensive representation of maize accessions and emphasizing landrace diversity. This diverse panel of domesticated maize and outgroups from across the Americas enables accurate genotyping in low-coverage samples while offering crucial context for interpreting diversity, particularly for natural diversity and paleogenomic analyses.

## Introduction

More than 10 years of whole genome short-read sequencing has yielded a wealth of genotypic data (Mardis, 2011) but integrating these legacy datasets is challenging due to highly heterogeneous coverage and read characteristics. Sequencing coverage in human genomic sequencing projects ranges from high (30-60X) (Bergström et al., 2020; Telenti et al., 2016) to moderate (4-20X) (1000 Genomes Project Consortium et al., 2015; Psaty et al., 2009) and even to skim sequencing (1-4X) (Rustagi et al., 2017); this is also the case in important crop species such as maize (Fig. 1A) (Bukowski et al., 2015; Gore et al., 2009; Hufford et al., 2021; Wang et al., 2017). Even with decreasing sequencing costs (Wetterstrand, 2022), low-coverage whole genome sequencing is a cost effective strategy to facilitate population-scale screening (Lou et al., 2021)(Parks & Lambert, 2015). The high diversity in both allelic and structural variation (Hufford et al., 2012) makes maize an ideal model to test the effects of coverage bias.

**Figure 1.**
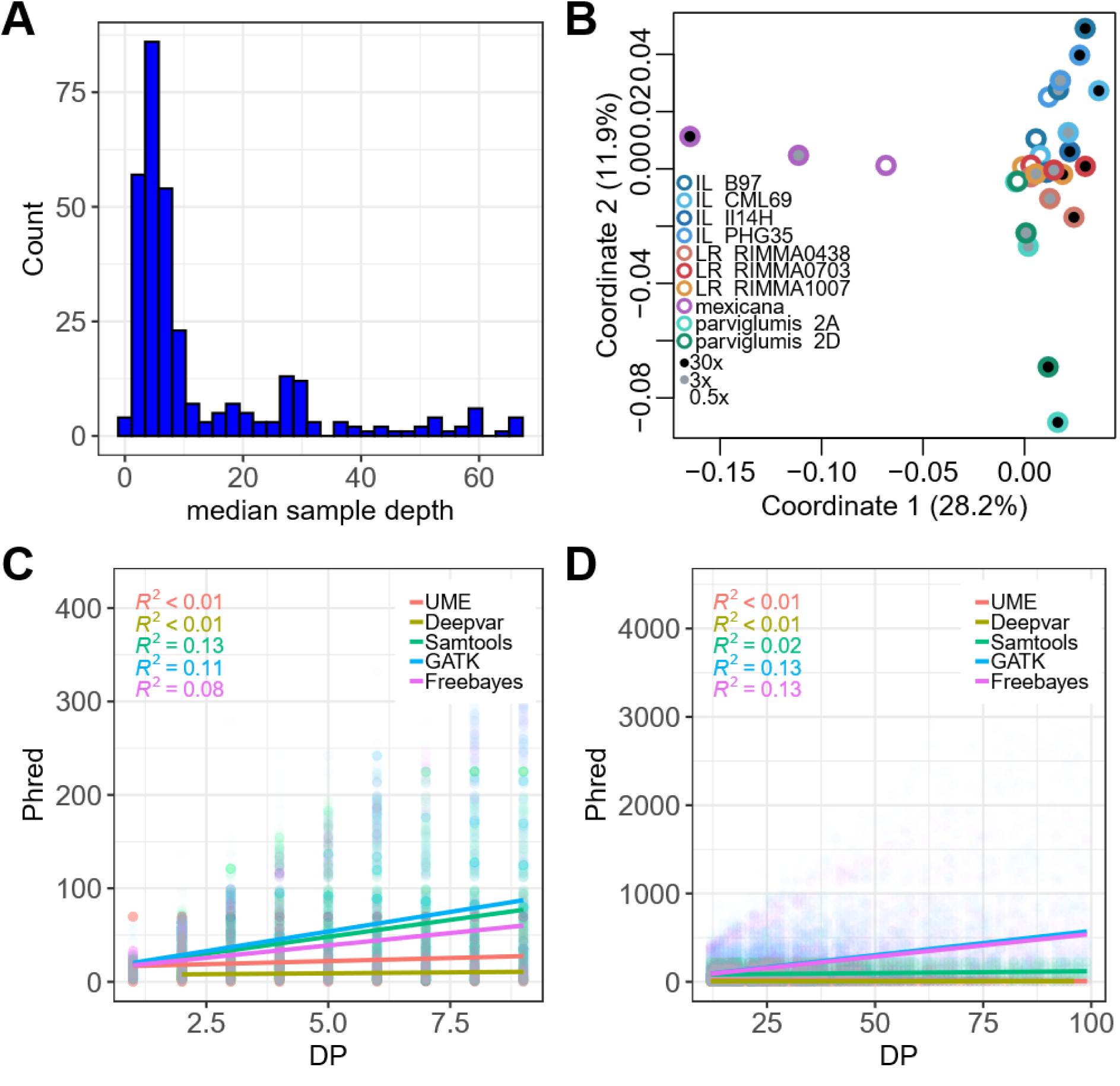
Effect of coverage bias over genotype likelihoods and genetic distances in standard callers. (**A**) Barplot showing the heterogeneity of depth coverage across public maize WGS samples. (**B**) MDS plot of downsampled maize accessions using the variant caller GATK. Border color indicates the identity of the sample, including inbred lines (IL), outbred landraces (LR), and maize progenitor *Zea mays* ssp. *mexicana* and *Z. mays* ssp. *parviglumis* subspecies. Fill color encodes average downsampled genome wide coverage (*SI Appendix*, Table S2). (**C and D**) Scatter plots of Phred-scale quality score versus depth (DP) for variants with 0.5-10X (C) and 11-100X (D) depth called using GATK, Samtools, Deepvar, Freebayes and the novel UME method. Colored lines correspond to the regression line of each caller (Materials and Methods).

Most whole genome sequence callers were developed for and optimized with high coverage libraries; as such, genotype calls in low coverage (<10X) samples can be biased towards the reference allele (Maruki & Lynch, 2017; Nielsen et al., 2012), impacting downstream diversity and population analyses (Benjelloun et al., 2019) for low coverage samples. This is particularly problematic when working with samples with heterogeneity in coverage because these differential biases can skew the estimated genetic relationships between individuals based on coverage (Fig. 1B,S4-S7) (Ross et al., 2013) and genetic distance from the reference genome. Additionally, while many high-quality sequence calling algorithms are available for high coverage samples (>30X) (Garrison & Marth, 2012; H. Li, 2011; McKenna et al., 2010), the choice of sequence caller can affect resulting genotype calls due to underlying differences in the algorithmic approach to error handling (Lin et al., 2022; Nielsen et al., 2012; Yao et al., 2020). Furthermore, the relationship between genotype accuracies and read depth is non-linear and relationships vary across callers (Fig. 1C and 1D).

Different strategies have been used to try to overcome such biases. For example, imputation of missing data using a reference panel (Ausmees et al., 2022; Y. Li et al., 2011; Rustagi et al., 2017; Swarts et al., 2014) can be very effective in panels with limited genetic diversity and extensive haplotype sharing with the reference panel. However, these are not accurate in diverse samples and novel variation will never be discovered. An intermediate approach to increase the number of genotype calls for low coverage samples is production calling, a method that uses a database of known trusted variant positions informed by multiple samples to report the SNP calls at trusted sites (Bukowski et al., 2018; Hui et al., 2020; Lou et al., 2021). The database of trusted variants requires a discovery step that includes *de novo* SNP calling and subsequent quality filters. Its utility for effectively calling variants in low coverage samples is a function of the genetic similarity between the individuals included in the discovery population and the target individuals.

The current work introduces an approach called **U**nified **M**ulti-caller **E**nsemble calibration (UME) to generate a high-quality discovery SNP database from heterogenous legacy data that can incorporate novel detected diversity from low and high coverage samples. The resulting variant set is suitable for production calling in diverse, low-coverage sequencing libraries from, for example, ancient or skim-sequenced modern individuals. We implement and optimize this approach in diverse inbred and outbred maize populations sequenced over the last decade with fold-coverages ranging from 1X to 90X per sample. Conceptually extending quality recalibration methods for low coverage sites (Chung & Chen, 2017; Le & Durbin, 2011), we accomplish this by employing an ensemble calibration approach that takes advantage of the strengths of callers built on diverse algorithmic approaches, normalizing and integrating genotype likelihoods before filtering individual genotype calls based on recalibrated likelihoods. These trusted variants can then be used in a production calling context to increase genotype calls for low-coverage samples (Fig. S1).

## New Approaches

In order to identify variants within individuals, standard callers such as GATK, Samtools, and Freebayes rely on combinations of different approaches (i.e. hidden Markov models, maximum likelihood, Bayesian inference) to define the error probability of a given variant. These approaches rely on repeated observations of a given variant to estimate error and therefore sites with higher coverage tend to have lower error probabilities. This association is even stronger for sites with less than 10X average coverage called with GATK, Freebayes and Samtools (Fig. 1C and 1D). The link between coverage and error probability has a profound impact on downstream analysis, as it affects the number of novel variants obtained and impacts the relationship between the target sample and the reference genome, ultimately affecting the relationships between analyzed samples and resulting in the observed coverage bias (Fig. 1B). On the contrary, the variant caller Deepvar does not model an error bias associated with depth (Fig. 1C-D) but rather with features extracted from a convolutional neural network. However, at low coverage (0.5 X), Deepvar reports 60% fewer variants than other callers (Fig. S2). Differences in the modeling of error probabilities or the quality of the variants resulting from each of the calling strategies lead to noticeable inconsistencies between the variants detected by the callers for the same sample (Fig. S3), which is a phenomena well described in the literature (Lin et al., 2022; Nielsen et al., 2012; Yao et al., 2020).

We propose a novel strategy to reduce the effect of coverage bias resulting from joint calling of heterogeneously covered samples by using an ensemble approach that uses genotype calls coming from orthogonal variant callers. This is attained by normalizing the different callers’ quality error estimates based on the empirical distributions of individual callers to make them comparable. The union of the recalibrated variants from the callers are merged as described below into a single summary statistic, integrating across one to four genotype calls. These error-prone genotype calls are subsequently filtered based on the joint summary statistic. Our tool performs the unified multi-caller ensemble calibration approach (UME) by taking advantage of the strengths of the different callers in order to extract as much information as possible across both low and high coverage samples. This approach is tested and optimized using publicly available maize samples (Table S1) that can be used later on for population genetics or association genomics. The main advantage of this approach is that it is population agnostic while still allowing representation of genotype calls from low coverage individuals in the final variant database. The resulting variant call set from the optimization and testing provides an important resource for the maize community as it represents a haplotype map of heterogeneous data treated under the same pipeline and corrected for coverage. Although the UME method has been tested on maize, it can be applied to datasets from other species.

### Discovery of trusted variants

The pipeline consists of two stages, discovery and production calling (Fig. S1). During the discovery call, all samples are independently called by each variant caller. The resulting Phred quality scores are assessed to generate an empirical cumulative distribution function of the modeled errors independently for each caller, thereby normalizing likelihood calls into a discrete distribution ranging from zero to one that can be directly compared across callers. (Fig. 2B and 2C). Then we perform a union (A ⋃ B) of all observed calls, based on preliminary testing (Fig. 2A, 2B and S8), coming from the different callers per position and per individual (missing genotype calls are included as ‘NN’). This standardizes the number of variants called for each individual. If observed alleles differ between callers, the genotype is generated from the two highest probability variants. The independently recalibrated quality scores from each caller (now scaled between 0 and 1, from most to least probable) can be combined in a number of ways into a summary statistic to determine the quality score for the merged variant calls; we tested the sum, max, min, mean and median of the recalibrated p-values for callers with non-missing variant calls. These error-prone variant calls are then filtered to produce a database of population-independent known variants. Filtering thresholds for the recalibrated variants are determined empirically for each discovery build based on maximizing the variance explained in the first four coordinates of metric multidimensional scaling decomposition of the pairwise genetic distance matrix. The approach minimizes error in the resulting set, as it is unlikely to be shared between individuals.

**Figure 2.**
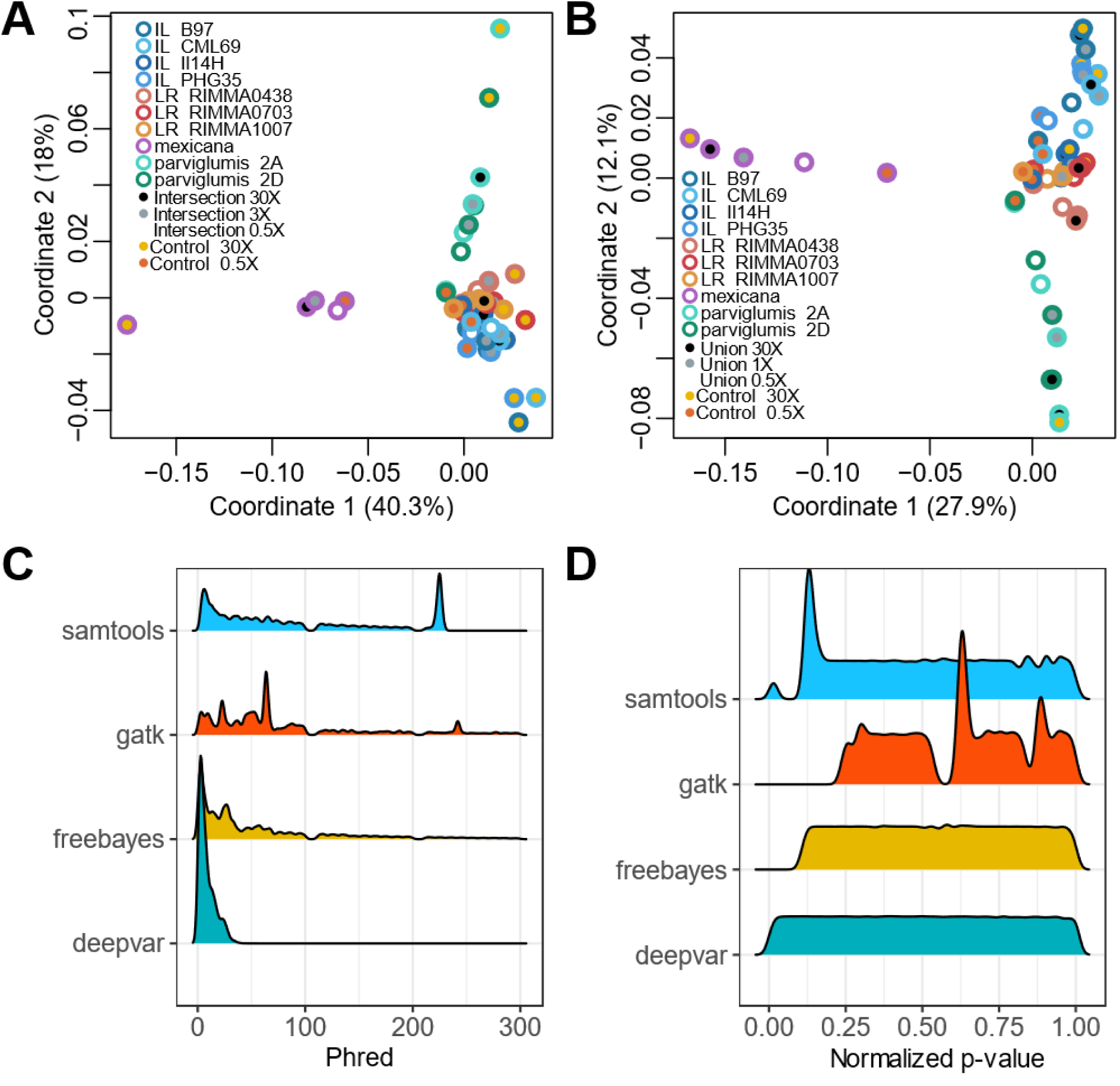
Effects of variant interaction and quality normalization. (**A**) MDS plot of downsampled maize accessions using the intersection between callers. Border color groups samples by inbred lines (IL), landraces (LR), and *mexicana* and *parviglumis* subspecies. Fill color encodes average downsampled genome-wide coverage at 30X, 3X, and 0.5X or control coverage at 0.5X and 30X (*SI Appendix*, Table S2). (**B**) MDS plot of downsampled maize accessions using the union between callers. Bolor color groups samples by inbred lines (IL), landraces (LR), and *mexicana* and *parviglumis* subspecies. Fill color encodes average downsampled genome-wide coverage at 30X, 3X, and 0.5X or control coverage at 0.5X and 30X (*SI Appendix*, Table S2). (**C**) Raw density distribution of quality Phred scores of standard variant callers. (**D**) Normalized density distribution of quality Phred scores of standard variant callers (see Materials and Methods).

### Production calling of variants from discovery call

The discovery phase results in a quality database of reliable variants but low coverage genotypes participating in the discovery build will still be under-called and potentially biased towards the reference genome because the low coverage alternate genotype calls are systematically biased against by the participating variant callers (Fig. 1). During the production calling phase, the dictionary of known variants per position is compared with mapped reads using mpileup (H. Li, 2011) and alleles matching a known variant are called even with only one mapped read in support. Finally, the variants are integrated into genotypes. If more than two variants are found in the sample for a given position, and those are present in the database, the two variants with the highest depths are kept as heterozygous. Paralogous positions were identified as those positions with more than one heterozygous genotype call in the inbred accessions and annotated in the VCF file as PARALOG on the info column. If a genotype is not in a position identified as paralogous, and the reads behind the two alleles are unbalanced as determined by a binomial approximation to a normal test for equality at two standard deviations, then it is set to missing.

### Validation and optimization

To explore the effect of coverage bias with respect to genetic diversity, we use several validation metrics on a coverage-controlled, downsampled dataset of 10 high coverage and diverse samples, the “*Downsampled diversity”* dataset (See Materials and Methods). Metric multidimensional scaling (MDS) plots are a means of visualizing genetic relatedness between individuals through the eigenvalue decomposition of a pairwise (Hamming) distance matrix. Similar to Principal Components Analysis, MDS is more robust against missing genotype data and in this dataset is particularly useful to visualize the effect of coverage bias between samples of different coverages. For the sample points in the MDS plots, the border color groups individual genotypes by inbred, landrace, mexicana, and parviglumis, while the fill color represents the downsampled coverage. If coverage bias is present in the samples then they will cluster by downsampling level rather than accession in MDS space. GATK exhibits a coverage bias that tends to cluster samples by coverage rather than by individual (Fig. 1B and Fig. S4). The same phenomenon occurs with other standard callers such as Samtools (Fig. S5), Freebayes (Fig. S6), and Deepvar (Fig. S7). Another validation metric is the comparison of the variants obtained between high coverage samples (See Materials and Methods, Table S2) from previous databases such as HapMap3 (Bukowski et al., 2018; Chia et al., 2012) and the variants obtained by this pipeline. Since in the high coverage samples we expect minimal bias due to coverage, site frequency spectrums and variance explained by MDS are useful indicators of biases associated with the proposed UME pipeline.

## Results

### Strategy for combining genotype calls across algorithms

We tested combining the genotype calls on the “*Downsampled diversity*” dataset as the intersection (A ∩ B) or the union (A ⋃ B) of the genotype calls from the four orthogonal variant callers (See Materials and Methods). The union produced 7-15% (along the 0.5-30x coverage range) more total positions than the intersection (Table 1 and Fig. S8). To test which method produced more biologically meaningful variation, we assessed genetic relationships using Metric MDS (R package cmdscale) on the IBS distance estimates between the downsampled individuals and those called with only low and high coverage GATK as a control (Fig. 2A and 2B). We observed that the intersection-treated samples tend to cluster with the low coverage controls independently from the simulated coverage, indicating a higher coverage bias than in the GATK-treated samples (Fig. 2A). On the contrary, for the union, we observed that samples clustered by accession rather than by coverage (Fig. 2B). For example, samples treated with the union 30X are closer to their corresponding 30X control GATK-treated samples (Fig. 2B). Interestingly, all 0.5X low coverage samples from the union pipeline cluster closer to the 30X than to the 0.5X low coverage GATK control (Fig. 2B). Because the union performed more similarly to the high coverage GATK control, we proceeded with the union of the positions.

**Table 1.** Mean count of detected SNPs by strategy and coverage for “*Downsampled diversity”* dataset.

| Coverage | Intersection | Union |
| --- | --- | --- |
| 0.5 X | 9,208,086 | 9,850,757 |
| 3 X | 11,539,048 | 12,934,711 |
| 30 X | 12,314,165 | 14,605,133 |

### Strategy for recalibrating error probabilities and filtering resulting genotypes

While the union of callers generates biologically meaningful variation independent of sample coverage, it also aggregates the error associated with each caller (Fig. 2C). We address this error by independently recalibrating the quality score (probability) distributions between 0 and 1 before integrating the normalized p-values into different summary statistics (max, min, mean, median) (Fig. 2D). These summary statistics yield different distributions of quality scores (Fig. S9). For example, the minimum error (min) reported by the union of calls generates the highest proportion of high Phred scores (low p-values) and is the most permissive calculation (only one caller needs confidence in the genotype), while the maximum p-value (max) has the lowest Phred scores of the participating callers, making it a more conservative estimator. The mean and median errors coming from the union of calls tend to be intermediate (Fig. S9).

We tested the impact of filtering based on the recalibrated summary statistics across a range of cutoffs to find the combination that generated the variant call set that best represented the genetic relationships between samples over the *“Inbred”* dataset (Table S2). To identify the recalibrated statistic and cutoff combination that explains most of the variance between samples, a proxy for genetically informative variation, we performed an MDS for each combination. As controls, we used both the high and low coverage GATK samples with the “*Downsampled diversity*” dataset, but also compared genetic relationships between another set of samples natively called on the *“NAM of HapMap3”* dataset (Bukowski et al., 2018; Chia et al., 2012) as well as the test here (Fig. S10).

To determine the optimum filtering criteria we assessed different filtered sets from the “*NAM of HapMap3*” build against native HapMap3 genotypes. We summarized the variance explained by the first four coordinate components for each MDS (Fig. 3B) as a proxy for identifying filter criteria that maximize shared genetic variation relative to unique and likely error-prone variant calls. The variance explained by the control HapMap3 equivalent variants is 65.2%; without filtering, the union explains only 61.9%. However, with filtering, an increase in variance explained is observed for all algorithms, for example, the maximum variance for the *min* is 69.7% at a normalized p-value cutoff of 8; in the case of the *median* the variance is 74.1% at a normalized p-value cutoff of 6; for the *mean* the variance is 75.6% at a normalized p-value cutoff of 8; finally the top maximum variance was observed when the *max* algorithm was applied at a normalized p-value cutoff of 5 with a 76% of the variance explained (Fig. 3B).

**Figure 3.**
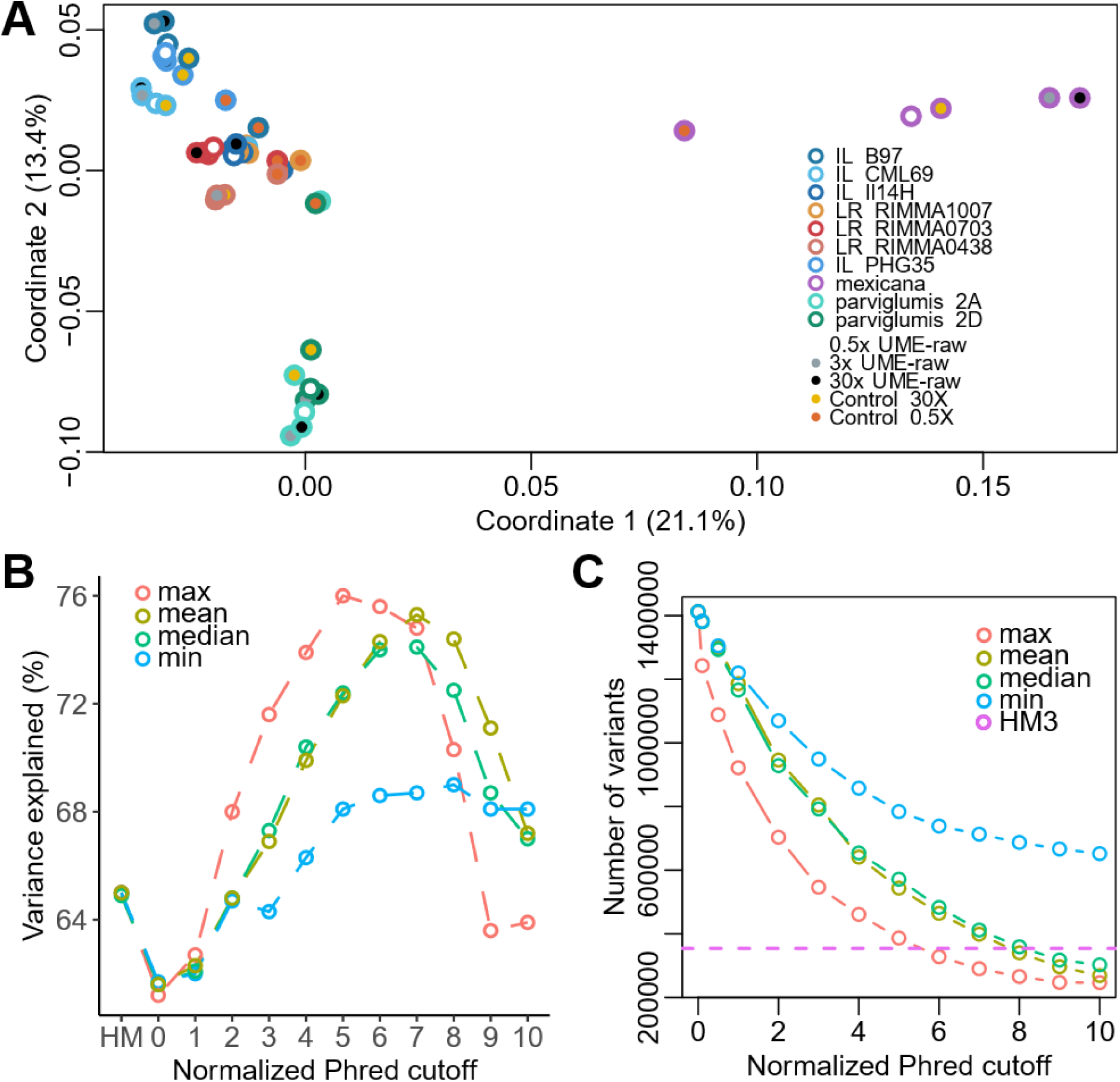
Parameter space exploration of quality scores in variant discovery. (**A**) MDS plot of downsampled maize accessions using the union between callers reporting maximum error and using a normalized p-value score cutoff of 5. Border color groups samples by inbred lines (IL), landraces (LR) and *mexicana* and *parviglumis* subspecies. Fill color encodes average downsampled genome-wide coverage at 30X, 3X, and 0.5X or control coverage at 0.5X and 30X (*SI Appendix*, Table S2). (**B**) Scatterplot of the variance explained in the first four components of MDS analysis over different normalized phred cutoffs and HapMap3 control (HM), comparing algorithms of union quality maximum, minimum, mean, and median (SI Appendix, Fig. S9). (**C**) Scatterplot of the number of variants obtained by applying different normalized phred cutoffs, comparing algorithms of union quality maximum, minimum, mean and median. Purple dotted line reflects the number of variants identified in the HapMap3 control.

This increase in variance for the max algorithm at the normalized p-value cutoff of 5 does not affect the overall relationships between the samples when compared to the control HapMap3 (Fig. S10 and Fig. S11), suggesting that these filtered sets maximize true genetic variance while minimizing variant call error. Additionally, the average IBS distance between each control HapMap3 accession and its corresponding UME-called sample (normalized p-value 5 cutoff) was 0.0089 (Fig. S11 and Fig. S12), which is below the average sequencing error. This filtered set shifts the site frequency spectrum towards a bimodal distribution of commonly shared and rare variants compared to the unfiltered union and the HapMap3 control (Fig. S13).

The unfiltered union had a 3.95 fold increase in the number of SNPs called compared to HapMap3; these are likely to be mainly spurious calls. *Min* tends to stabilize at 2.25 fold. *Mean* and *median* reach the control amount of variables at a cutoff of 8 and for the max algorithm the cutoff of 5 represents a fold change increase of 0.09 (Fig. 3C).

### Impact of filtering criteria on the “Downsampled diversity” dataset

We defined the max at a normalized p-value cutoff of 5 for downstream analysis, as described previously, and the resulting filtered call set constitutes the *de novo* Discovery build. We tested the resulting call set with respect to the *“Downsampled diversity”* dataset to understand how the filters affected coverage bias. In MDS space (Fig. 3A), filtered individuals cluster by sample rather than by coverage and also cluster with the control samples (the same genotypes at high coverage called with the default GATK pipeline), suggesting that the coverage bias was corrected by the union max strategy. UME uncoupled coverage and the estimated error rate, as the adjusted R^2^ for depth versus error was less than 0.006 for all UME strategies (Fig. S14). Using a cutoff of 5 further broke this relationship between error and coverage, with an R^2^ of 0.01 (Fig. S15). This pattern is the same when looking at the sites with a depth lower than 10X, where the R^2^ is lower than 0.01 (Fig. 1C).

### Production calling increases the proportion of genotype calls for low coverage individuals

We re-called all samples from the mapped bam files at only the positions and for alleles discovered in the previous step as a final “production call” in order to increase genotype calls for low-coverage samples. When comparing *Discovery* and *Production* results we identify a decrease in the proportion missing at sites present in the *Production* stage (Fig. 4B). The reduction of the proportion missing affects all coverages from 0.5X to 30X; however, the effect is maximized in the low coverage samples (Fig. 4C). In the “*Downsampled diversity”* dataset, we observe a clustering of samples by accession rather than by coverage (Fig. 3A and 4A).

**Figure 4.**
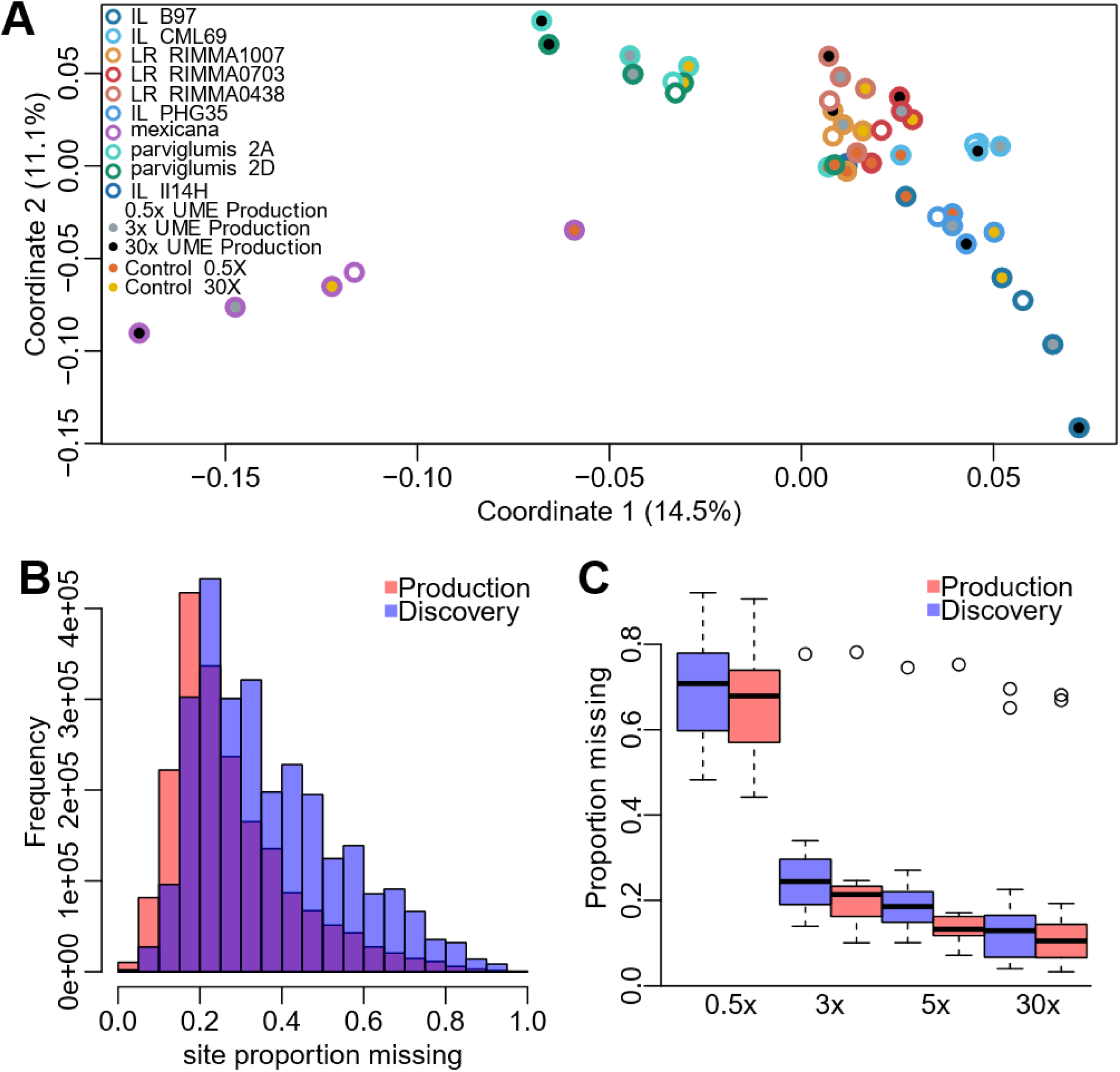
Effect of production call over coverage and genetic distances. (**A**) MDS plot of downsampled maize accessions using variants generated by production calling with the union max 5 cutoff. Border color groups samples by inbred lines (IL), landraces (LR), and *mexicana* and *parviglumis* subspecies. Fill color encodes average downsampled genome-wide coverage at 30X, 3X, and 0.5X and control coverage at 0.5X and 30X (*SI Appendix*, Table S2). (**B**) Histograms of site proportion missing comparing discovery and production calling of Union max 5 cutoff. (**C**) Box plots of site proportion missing by the different taxon coverages at 0.5X, 3X, 5X, and 30X comparing discovery and production calling of union max 5 cutoff.

### UME for calling genotypes for a diverse maize HapMap database

A new maize diversity build is needed for two main reasons. As we and others have shown (Ross et al., 2013), differential coverage can generate biased genetic relationships when using standard callers with default parameters. To overcome this, some previous builds (Bukowski et al., 2015; Chia et al., 2012; Gore et al., 2009) have used custom population filters, but this can result in ascertainment bias where more frequently represented haplotypes are overrepresented in the resulting SNP set. UME is resistant to both of these biases. Secondly, the diversity represented within previous variant sets is imbalanced between accessions over geography, improvement history and between in– and out-groups (Table. 2). Outgroups and landraces contain more diversity than is present within the inbred populations (Romero Navarro et al., 2017; Wang et al., 2017) and when they are underrepresented in variant discovery populations, this variation is often lost, especially in low-coverage samples filtered using population metrics. This diversity is useful for exploring evolution and adaptation and can better contextualize new diversity (Vielle-Calzada et al., 2009), including archaeological maize samples. The build presented here consists of 818 accessions, including 188 inbred lines, 356 landraces, 95 *Zea mays* ssp*. parviglumis*, 84 *Zea mays* ssp*. mexicana*, 88 other *Zea* outgroups such as *Zea luxurians*, *Zea nicaraguensis*, *Zea perennis*, *Zea diploperennis*, *Zea mays* ssp*. huehuetenangensis* as well as five distant *Tripsacum dactyloides* outgroups (Table. 2 and Table S1). Data was obtained from public sources (Chia et al., 2012; Gault et al., 2018), including data from the following SRA Bioprojects: PRJEB31061 (Hufford et al., 2021), PRJNA300309 (Wang et al., 2017), PRJNA381642 (J. Yang et al., 2017), PRJNA389800 (Bukowski et al., 2018), PRJNA479960 (Kistler et al., 2018), PRJNA641489 (Chen et al., 2022), PRJNA783885 (N. Yang et al., 2023).

**Table 2.** Comparison between maize HapMap diversity panels.

| Source | Chia et al.<br>2012 | Bukowski et al.<br>2018 | Grzybowski et al<br>2023 | This study |
| --- | --- | --- | --- | --- |
| Number of SNPs | 55,061,920 | 83,153,144 | 43,296,332 | 64,531,167 |
| Total accessions | 103 | 1218 | 1515 | 818 |
| Inbred lines | 60 | 1173 | 978 | 189 |
| Landraces | 23 | 23 | 142 | 356 |
| *Zea mays* ssp.<br>*parviglumis* | 19 | 19 | 88 | 95 |
| *Zea mays* ssp. *mexicana* | 2 | 2 | 79 | 84 |
| Other *Zea* ancestors | 0 | 0 | 54 | 88 |
| *Tripsacum dactyloides* | 1 | 1 | 1 | 5 |

Based on the parameters identified in the *Downsampled diversity* dataset, the full maize variant call set was then built using UME Discovery by calling variants over the previously described 818 accessions, using the union of four different callers (GATK, Samtools, Freebayes and DeepVar; see Materials and methods) filtered with the max error summary statistic. To identify the proper cutoff for the error probabilities, we randomly subsampled 3 million variants using various cutoffs from 1 to 20. Each resulting discovery set was production called in order to identify the cutoff that maximizes variation underlying genetic relationships between individuals while minimizing error (Fig. S16). We identified that the cutoff of 10 explained the most variance in MDS with a total of 12.8% over the first four components. This step of recalibrating the error cutoff is crucial for every discovery dataset, as the error rate is determined by the variation within the population. Using a cutoff of 10 over the whole dataset, we identified a total of 64,531,167 SNPs; INDELS were not included due to difficulties in modeling the probabilities over infinite states.

To evaluate the magnitude of spurious variants over the “*Inbred”* dataset we used the 22 inbred lines that have been subjected to at least 5 rounds of selfing (NAM) and that have been independently sampled four times from the same accession in the dataset. In inbred lines, heterozygosity results from genotyping error, paralogy, or residual heterozygosity. Within the isolated 3 million positions (see Materials and methods), we identify that, on average, 0.36% of the sites are heterozygous singletons, i.e. present only within one replicated inbred, thus establishing a maximum residual sequencing error. This may be an overestimate as some might still be low frequency paralogous genotypes, however, as singleton heterozygosity increases with high coverage where there is power to detect rare variants and a certain number of new mutations are expected in an individual (Fig. S17). To better focus on true genotyping error, we evaluated the proportion of singletons that showed allelic imbalance at more than two standard errors (based on the binomial approximation to a normal distribution), which reduced the estimated error (at these heterozygous positions) to 0.12%. These positions were then filtered from the dataset (a total of 89,126 positions). The genotypes that were identified as unbalanced singletons were set to missing in the final dataset. To identify paralogous positions, we focus only on genotypes that have at least 9 reads in order to accurately estimate the heterozygous frequency at a site. Conservatively, any position with at least two heterozygous genotype calls is annotated as a PARALOG (43,908,437 positions, 68% of all positions). Interestingly, a subset (20.89%) of the paralog-annotated positions are overrepresented with unbalanced heterozygosity, suggesting tandem duplications or other more complicated structural variations (Fig. S18).

We further tested the similarity of independently sequenced inbred genotypes in our database derived from different experimental sources, such as the maize 282 association panel (Bukowski et al., 2018) with an average coverage of 4X, HapMap2 (Chia et al., 2012) with an average coverage of 2X, HapMap3 (Bukowski et al., 2018) with an average coverage of 5X, and the NAM *de novo* assembly (Hufford et al., 2021) with an average coverage of 40X (Table S1). We identified that the samples group by accession independently from the source or coverage in MDS space (Fig. S19), suggesting that the UME strategy successfully overcame coverage bias in our maize haplotype database. We applied our coverage bias corrected maize haplotype database across the *Zea* genus and *Tripsacum,* and the first two coordinates of the MDS cumulatively explained 24.1% of the total variance (Fig. 5A). The first two coordinates of MDS within domesticated maize only reflect the 3 main edge populations from US, Mexico, and South America, including a high-density sampling of admixed individuals along these edge populations (Fig. 5B). The folded site frequency spectrum of our HapMap suggests that most of the variance is shared between samples with an average MAF of 0.09. We were able to confidently call singletons relatively rarely; only 0.46% of the variants called are singletons (Fig. 5C). Variant sites are covered at a mean proportion of 24% of the individuals with a minimum of 4% (Fig. 5D) and a mean heterozygosity of 14% of the individuals per site (Fig. 5E). Since we have accessions that are inbred lines with several rounds of selfing, as well as open pollinated landraces and teosintes (Table S1), the distribution of heterozygosity across taxa is multimodal (Fig. 5F). Finally, the database has differences across proportion missing per taxon, where the mean proportion missing is about 24.34% (Fig. 5G and Table S1), likely attributable to the structural variation present in maize and consistent with previous maize diversity builds (Bukowski et al., 2018; Chia et al., 2012; Grzybowski et al., 2023).

**Figure 5.**
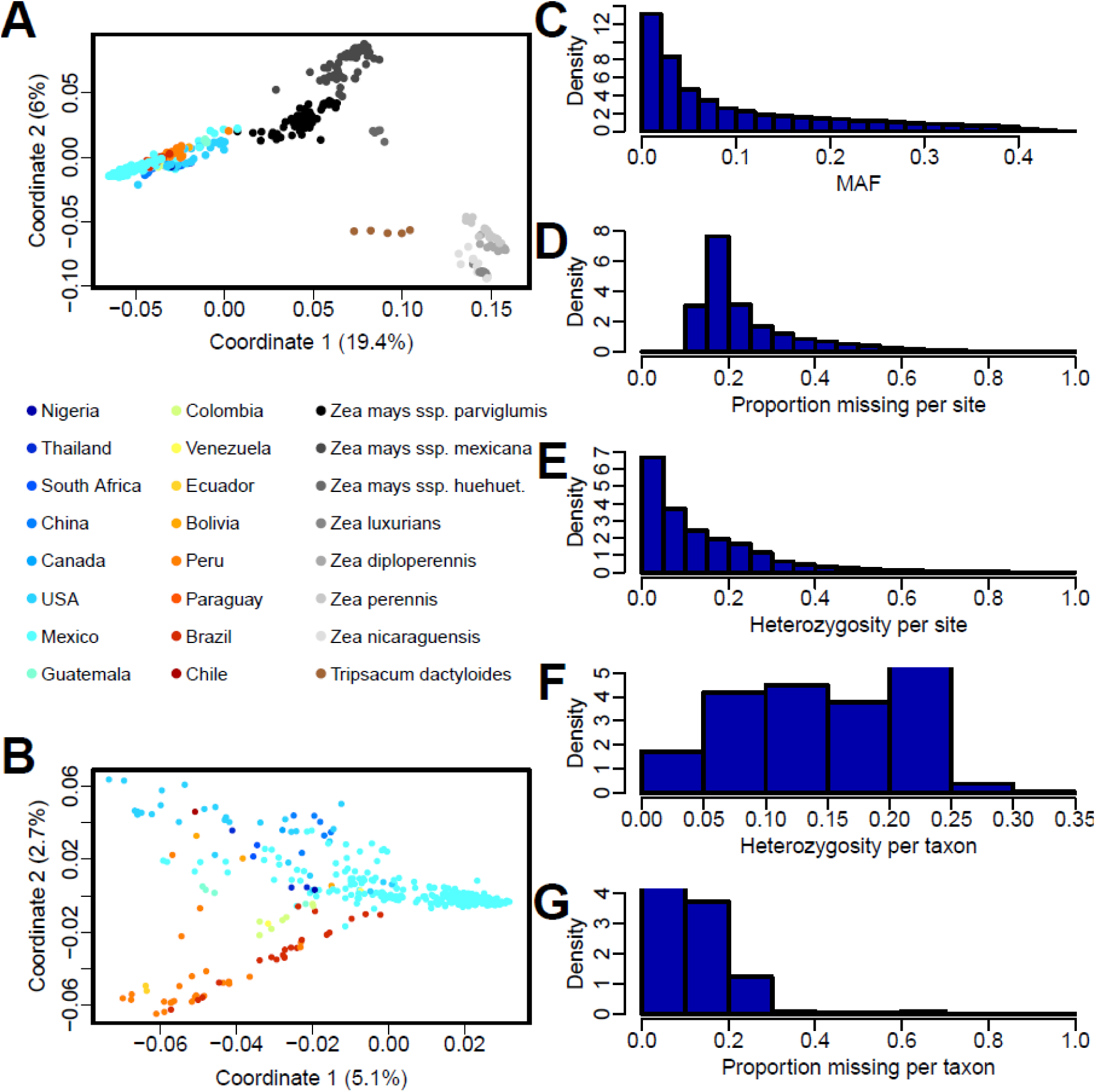
Metrics of coverage controlled maize database. (**A**) MDS plot of the diverse accessions in the database including close and distant ancestors. Domesticated *Zea mays* ssp*. mays* accessions are colored by country of origin, *Zea mays* wild relatives are colored in grayscale, and the distant ancestor *Tripsacum dactyloides* is colored in brown. In the legend, *Zea mays* ssp. *huehuetenanhensis* was shortened to *Zea mays* ssp. *huehuet*. (**B**) MDS plot of *Zea mays* ssp*. mays* accessions in the database without wild relatives. Accessions are colored by country as in (A). (**C**) Folded site frequency spectrum of the resulting SNPs, where the X-axis shows the Minimum Allele Frequency (MAF) and the Y-axis is the relative density per MAF value. (**D**) Site coverage profile of the SNP dataset, where the X-axis shows the proportion of missing alleles between the individuals for each site and the Y-axis the relative density. (**E**) Heterozygosity profile of the SNP dataset, where the X-axis shows the proportion of heterozygous individuals for each site and the Y-axis the relative density. (**F**) Heterozygosity profile for each individual in the dataset, where the X-axis shows the proportion of heterozygous SNPs for every taxon and the Y-axis the relative density. (**G**) Coverage profile for each individual in the dataset, where the X-axis shows the proportion of missing alleles SNPs for every taxon and the Y-axis the relative density.

## Conclusion

Variant calling in low-coverage data remains a challenge, especially for population analyses where coverage and ascertainment bias represent a major source of error downstream. We have provided evidence that the combination of calls from orthogonal variant callers outperforms individual strategies during *de novo* calling in mixed-coverage datasets. Our Unified Multi-Caller Ensemble (UME) approach achieves variant calling in data with coverage as low as 0.5X while preserving the biological information of the resulting variants. Quality recalibration of the identified variants is a core stage of the process, enabling the direct comparison of probabilities between callers while breaking the correlation between quality and coverage. Based on the evidence presented, we conclude that using the union of calls with a conservative error adjustment that returns the maximum p-value across callers followed by biologically informed filtering of the discovery dataset based on maximizing shared variance and subsequent production calling generates a reliable call set that minimizes error and coverage bias, retains significant structural variation as paralogous positions and retains variation from genotypes distantly related to the reference. One significant benefit of this approach is that it is not influenced by the frequency of variants in the discovery panel, which can be biased by the population structure of the individuals used. Even with the heterogeneity of accessions in the “*Downsampled diversity”* dataset, genetic distances were consistently improved and matched the high-coverage equivalent controls. Therefore, UME can be considered a population-agnostic variant calling strategy.

Our results demonstrate that the production stage enriches the identification of variants at all coverage levels, particularly in low-coverage samples, without sacrificing accuracy, as would be the case with conventional callers. We present a coverage bias-controlled HapMap database for a balanced set of maize accessions, representing an up-to-date sampling of maize diversity that favors landrace diversity. This diversity panel empowers low-scale genomic analysis of specific accessions by providing a baseline framework, allowing accurate variant calling for low-coverage samples using the UME-Production call. Simultaneously, it contextualizes the diversity obtained from those low-coverage samples, particularly in paleogenomic analysis where sample variance arises from both low-coverage data and randomly distributed molecular damage across the genome. Our presented HapMap build has the unique feature of not being filtered by linkage disequilibrium, allowing detection of diversity in low-coverage samples below 1X. Instead of aiming to find a single variant over a recombination block, which is unlikely due to low coverage, it detects other variants within the same recombination block, providing a better framework for genome-wide diversity patterns. Additionally, the inclusion of the dataset of Mexican landrace accessions (N. Yang et al., 2023) increases the representation of the diversity close to the center of origin, balancing the variance of the discovery dataset. The balanced HapMap presented here provides a context for investigating evolutionary questions in maize across the Americas.

## Materials and Methods

### Datasets

Three populations were used for testing and validation, hereafter referred to as the “*Downsampled diversity*”, “*NAM of HapMap3*” and “*Inbred*” datasets. To explore the effect of the interaction between variant information coming from orthogonal callers, we used a limited, high diversity, high coverage set of individuals (“*Downsampled diversity*” panel), including previously published genomic data of 10 accessions (Table S2) from the *Zea* genus: five inbred lines (*Zea mays* ssp. *mays*), three landraces (*Zea mays* ssp. *mays*), two teosinte *parviglumis* (*Zea mays* ssp. *parviglumis*) and one mexicana (*Zea mays* ssp. *mexicana*) individual. Samples were nested downsampled to 30X, 3X, and 0.5X using the GATK v 4.2.0 (McKenna et al., 2010) downsample function and checked for average coverage using the Samtools (H. Li, 2011) coverage function. This dataset consists of a total of 18,655,847 SNPs within the full alignment. As a control of a standardized pipeline we included equivalent individuals called only by GATK at 30X and 0.5X simulated coverages (Fig. 2A and 2B).

The second dataset, “*NAM of HapMap3***”,** allowed a validation of UME parameters by comparison with the previous public diversity panel (Bukowski et al., 2018), as well as determination of parameters. The panel includes 17 high-coverage maize inbred lines with an average coverage of 30.8X (Table S2) (Bukowski et al., 2018).

The third dataset, “*Inbred*”, consists of a set of 22 inbred lines that have individuals from the same accession coming from four different sources, the 282 maize diversity panel (Bukowski et al., 2018), HapMap2 (Chia et al., 2012), HapMap3 (Bukowski et al., 2018), and NAM *de novo* sequencing individuals (Hufford et al., 2021). This panel allowed us to estimate the residual error and to annotate the paralogous sites.

The last dataset is the resource generated “*Zea diversity panel*’’ and comprises 818 accessions (Table S1) that include the raw reads from public sources (Chia et al., 2012; Gault et al., 2018) including data under the Bioprojects: PRJEB31061 (Hufford et al., 2021), PRJNA300309 (Wang et al., 2017), PRJNA381642 (J. Yang et al., 2017), PRJNA389800 (Bukowski et al., 2018), PRJNA479960 (Kistler et al., 2018), PRJNA641489 (Chen et al., 2022), PRJNA783885 (N. Yang et al., 2023). The HapMap presented here can be publicly accessed upon final publication in vcf format.

### Variant calling pipeline

All samples were mapped to the current maize reference genome B73_V5 (Hufford et al., 2021) using BWA-MEM v 0.7.17 (H. Li & Durbin, 2010). Variants were called with independent samples using GATK v 4.2.0 (McKenna et al., 2010), freebayes version v 1.3.5 (Garrison & Marth, 2012), Samtools v 1.10, bcftools mpileup v 1.9 (H. Li, 2011), and nf-core/deepvar v 1.0 (Cheng et al., 2020). Raw vcf files were used for downstream analysis. The control SNP dataset included equivalent genotypes from the previously published 282 maize diversity panel in HapMap3 (Bukowski et al., 2018; Chia et al., 2012). Hapmap3 was natively called under refgen v4 and the variants here were converted with crossmap v 0.5.1(Zhao et al., 2014) to the current maize reference B73_V5 (Zm-B73-REFERENCE-NAM-5.0) (Hufford et al., 2021) downloaded from maizeGDB (Woodhouse et al., 2021). IBS distance matrices, SFS, site proportion missing and taxon coverage were produced using TASSEL 5.2.68 (Bradbury et al., 2007). All plots including histograms, box plots, distributions, bar plots, scatter plots, linear regressions and multidimensional scaling (MDS) were produced with cmdscale stats version 3.6.2 in R v 4.1.2 (Team, n.d.).

### Sampling SNP set for testing

For the *“Inbred”* and “*Downsampled diversity”* datasets, 10,000,000 SNPs were randomly sampled from all sites present in chromosome 10. From those sites, quantifications of the number of variants or polymorphic sites were reported in downstream analysis. For example, 354,142 variants were identified in the HapMap3 controls.

### Strategy for combining genotype calls across algorithms

We report two different strategies of combining genotype calls across algorithms. First we tested the intersection (A ∩ B) of variants between callers. Therefore, positions per individual were kept if that position matched the same variant between all four callers. The second strategy tested was the union (A ⋃ B) of variants between callers. For this strategy all variants per position and individual were kept; if genotype calls differed between callers, the two most likely ones were kept as candidates for that position as potential alleles. During the “discovery” stage a list of potential alleles observed in all individuals per position was created. This variant database is used in the “*production*” stage to call genotypes for each individual constrained by variants present in the database. If two or more variants are identified in an individual, the most likely ones based on the calibrated probabilities are kept as heterozygous. All missing variants are reported within the alignments and kept as “NN”.

### Residual error and paralog annotation

In order to determine the amount of residual error in the dataset as well as to annotate the paralog positions in the panel, we used the “*Inbred*” dataset. We sampled 3 million sites randomly from the “*Zea diversity panel*” for those individuals. All individuals chosen are inbred lines that were NAM founders and therefore low levels of heterozygosity are expected. Meanwhile, shared heterozygous genotypes are likely to be real and part of the structural variation or heterozygotes maintained by selection, while heterozygous singletons are likely to be spurious variants and part of the residual error of the dataset. To estimate the residual error we quantified the amount of singleton heterozygous positions and divided them by the total 3 million sampled positions. For those singleton heterozygous positions, we tested if the alleles A and B were balanced or not. The expectation in a diploid is that real heterozygous positions will tend to appear as balanced alleles due random sampling. As such, we quantified the amount of singleton heterozygous positions that were unbalanced under the two sigma probability based on the binomial approximation to a normal and then divided the sum of those positions over the total sampled sites and reported that value as residual error of the dataset. To annotate the paralogous positions we focused on genotypes that had at least nine reads for the genotype, giving less than one percent chance that a heterozygote will be under called as homozygous. We calculated the two sigma probability based on the binomial approximation to a normal of that genotype to be unbalanced, meaning that allele A and B of the heterozygous genotype did not have the same ratio. If the genotype is an unbalanced singleton heterozygous position, then it is set to missing. Therefore, unbalanced heterozygous genotypes that are not singletons are likely to be paralogs. Positions with unbalanced heterozygous genotypes were annotated with the legend “PARALOG” at the given position in the INFO column of the vcf file.

### UME usage

UME is written in Perl v 5.32.0 and deposited in a public repository (https://github.com/Vallebueno/UME). UME has three main functions: 1) *Normalize*, which transforms the Phred scores into the normalized distribution based on the provided empirical distribution, 2) *Discovery*, which applies a cutoff to the normalized Phred scores to generate the list of trusted alternative alleles for each position of the genome, and 3) *Production*, which calls the variants of individual samples from the read alignment based on the trusted variants found in the discovery step. The input of the discovery function includes a multi-vcf file with the variants identified by publicly available callers and a key file with the information on the localization of each sample in the multi-vcf and the name of the caller for each column of variants. Therefore the number of columns of the file should be the number of individuals times the number of callers used. The algorithm creates the following outputs: a multi-vcf file with the number of columns equal to the number of individuals, and a dictionary of known variants for each position based on the individuals used in the original input. User-modified parameters available include: ‘-a’, which allows one to use a different method of interacting with the error probabilities (max (default), mean, median, min, sum), and ‘-c’ that allows one to impose a filter based on the final empirical probabilities (0-9). This value is dependent on the structure of the data and has to be tested looking at the variance in MDS space for the user dataset. The input of the production function is the two files resulting from the discovery step plus the read information file (mpileup) of each sample. The two functions are kept independent to allow for parallelization.

## Supporting information

Annex Data

## Acknowledgments

We thank Laura Morales and Haijun Liu, for giving constructive feedback about the manuscript. This project has received funding from the European Union’s Framework Programme for Research and Innovation Horizon 2020 (2014-2020) under the Marie Curie Skłodowska Grant Agreement Nr. 847548

## Supplementary Materials

**Figure S1.**
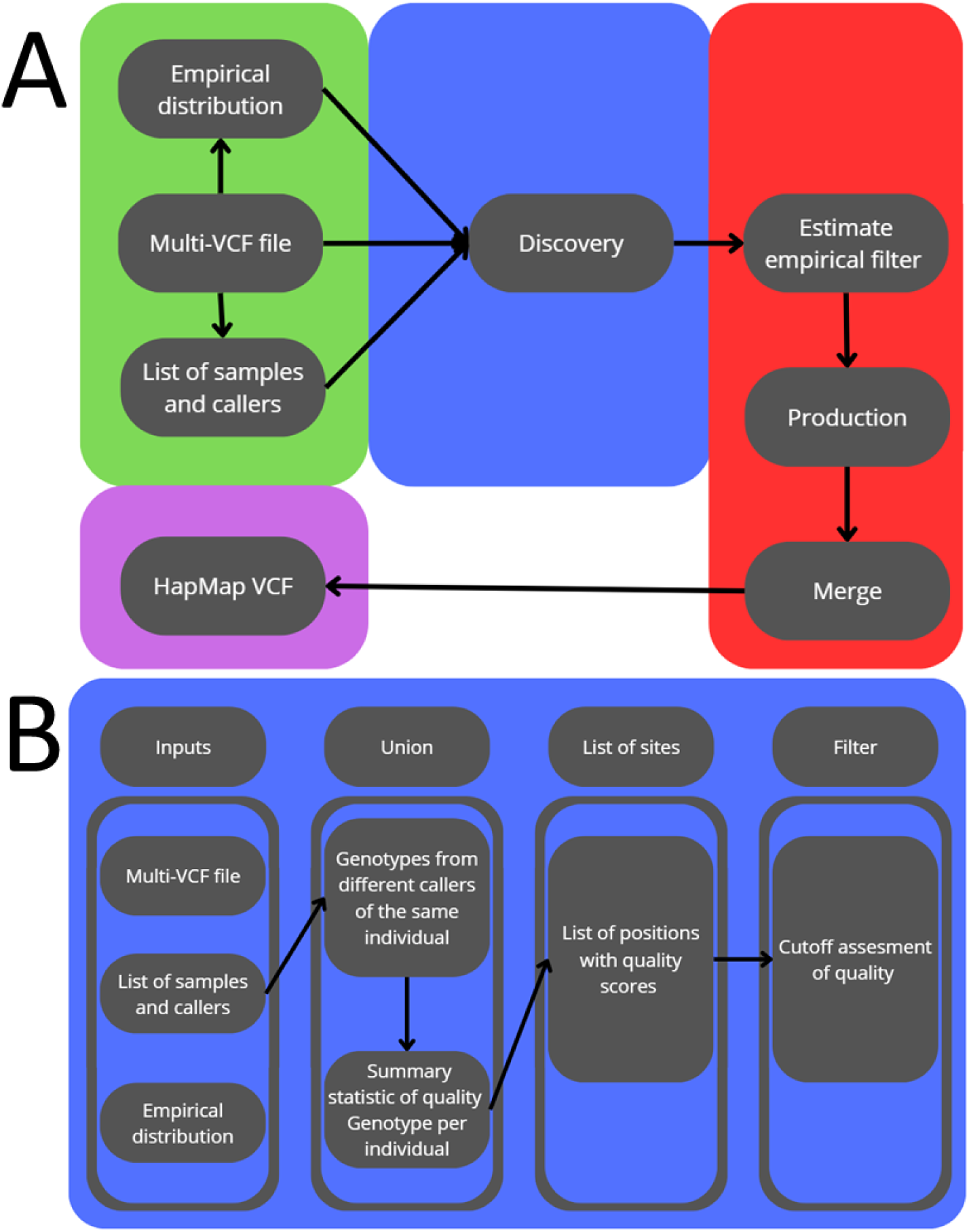
– Flux Diagram of the UME Pipeline. **A**) Within the UME pipeline, a well-defined flow of operations emerges, represented through a comprehensive flux diagram. This pipeline revolves around three core functions, each color-coded for clarity and cohesion. Clist Function (Green): At the heart of the pipeline, the Clist function takes a multivcf as input and initiates the process by creating empirical distributions of phred scores for each of the callers. This step forms the foundational basis for subsequent analyses. Discovery Function (Blue): The Discovery function plays a pivotal role by generating a list of trusted alleles per position. This process is governed by strict criteria, including the determination of maximum error and an assigned cutoff. The results are used to filter and extract the most reliable data. Production Function (Red): The Production function leverages the empirical filter calculated by the user. This filter is often based on a comprehensive variance analysis, as exemplified in figure S16. Using mpileup files and the Discovery list of trusted alleles, the Production function generates individual files for each sample. These individual files are later merged through a bash command, culminating in the creation of the coveted HapMap VCF file. B) Flux diagram of the discovery function of UME

**Figure S2.**
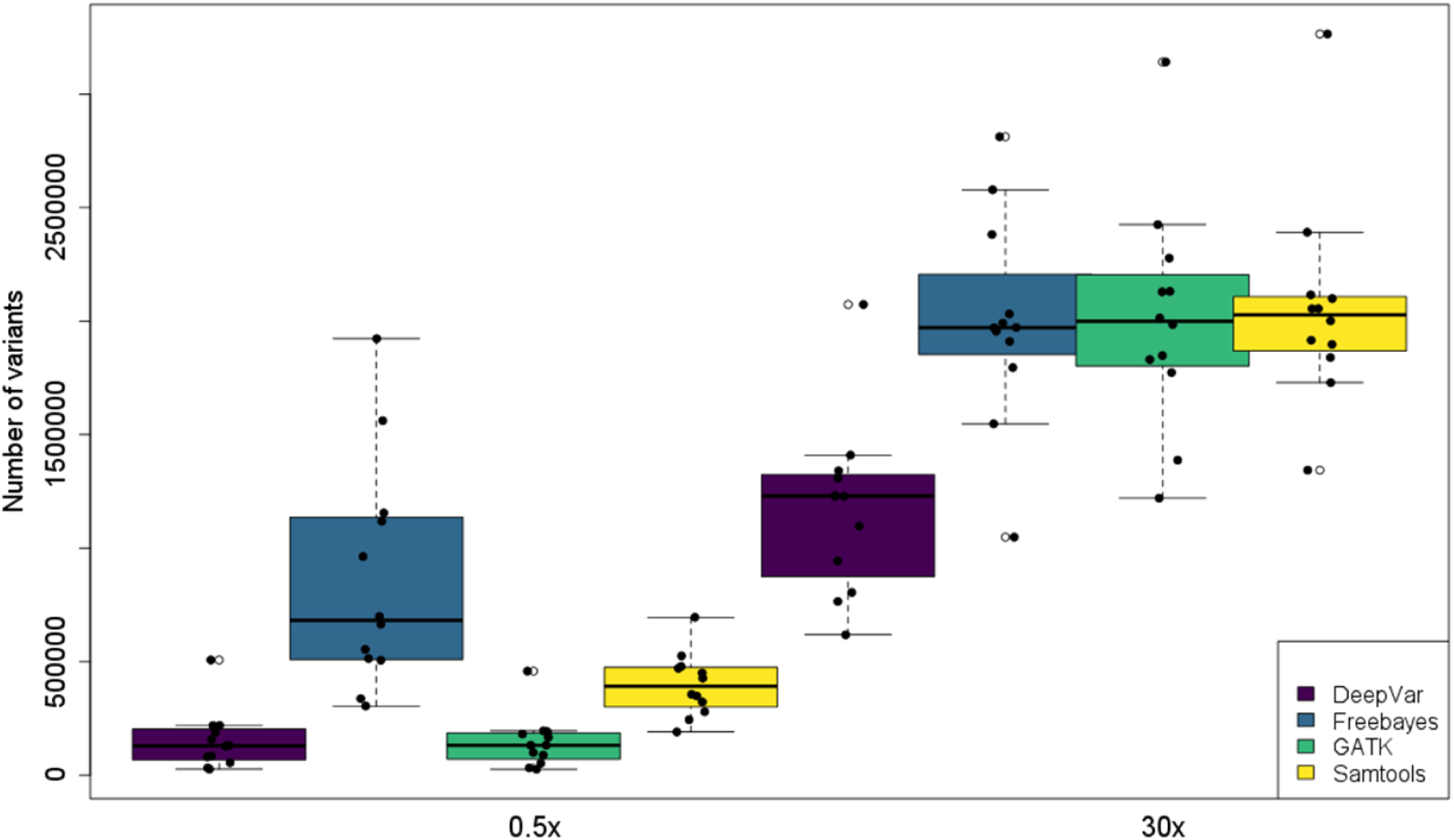
– Quantification of raw variant detection across multiple variant callers at varying sequencing depths. The Y-axis represents the total count of identified variants with a Phred quality score exceeding 0, derived from an analysis of downsampled samples (*SI Appendix*, Table S2). Different variant callers are distinguished by distinct colors. The X-axis displays data grouped into blocks of four callers, representing low sequencing coverage (0.5X) and high sequencing coverage (30X).

**Figure S3.**
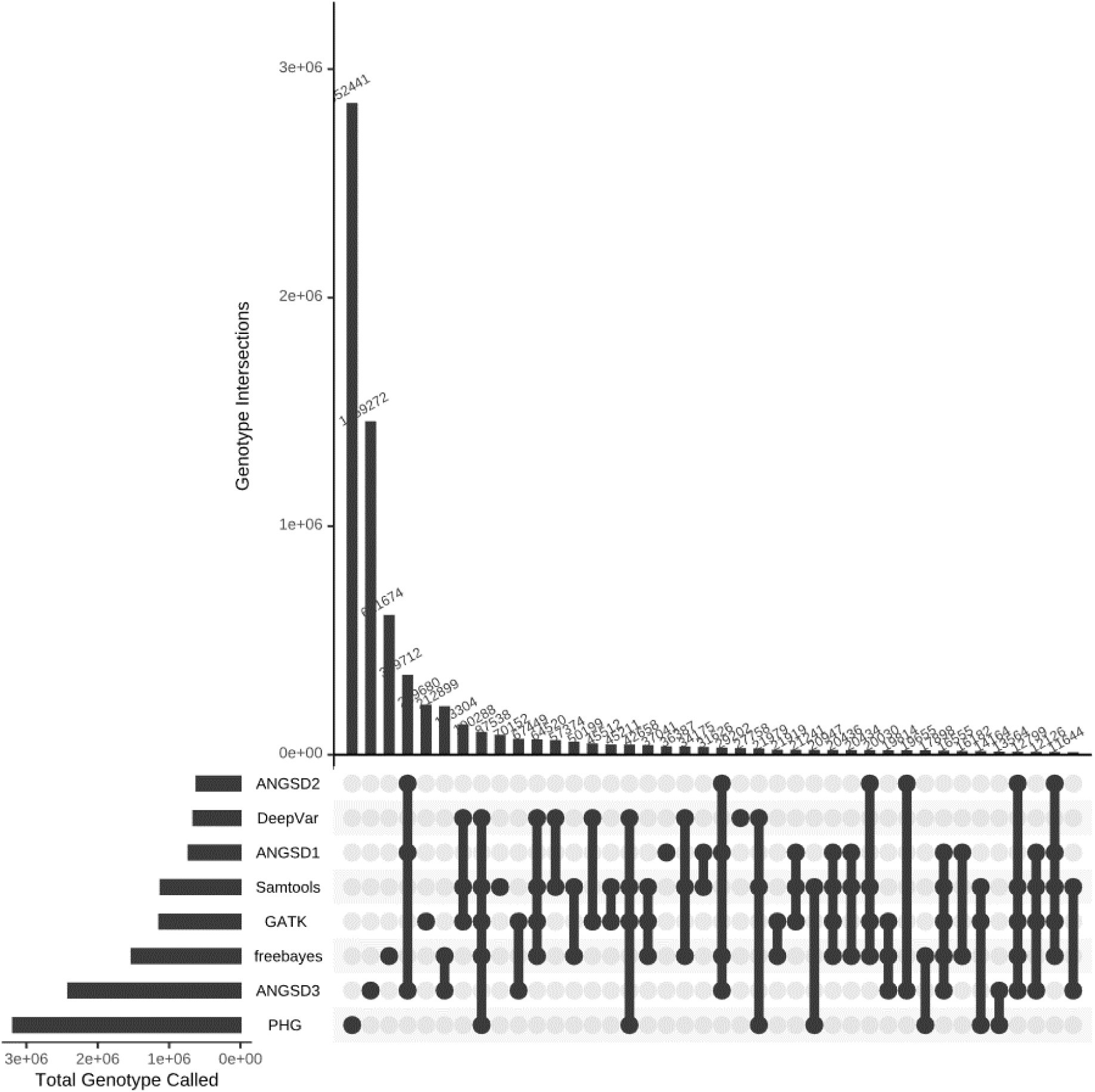
– Quantification of variant consistency among different callers within a single individual. The UpSetR Plot illustrates the intersection of genotype frequencies for the B97 inbred line. Each column represents a unique combination of variant callers, while each row corresponds to a specific caller. Shared intersections of genotype frequencies among caller combinations are indicated by dots aligned within the columns. The horizontal bars positioned at the top of the plot provide an overview of total genotype frequencies for each individual caller, whereas the vertical bars on the right-hand side represent the cumulative frequency of each genotype across all callers.

**Figure S4.**
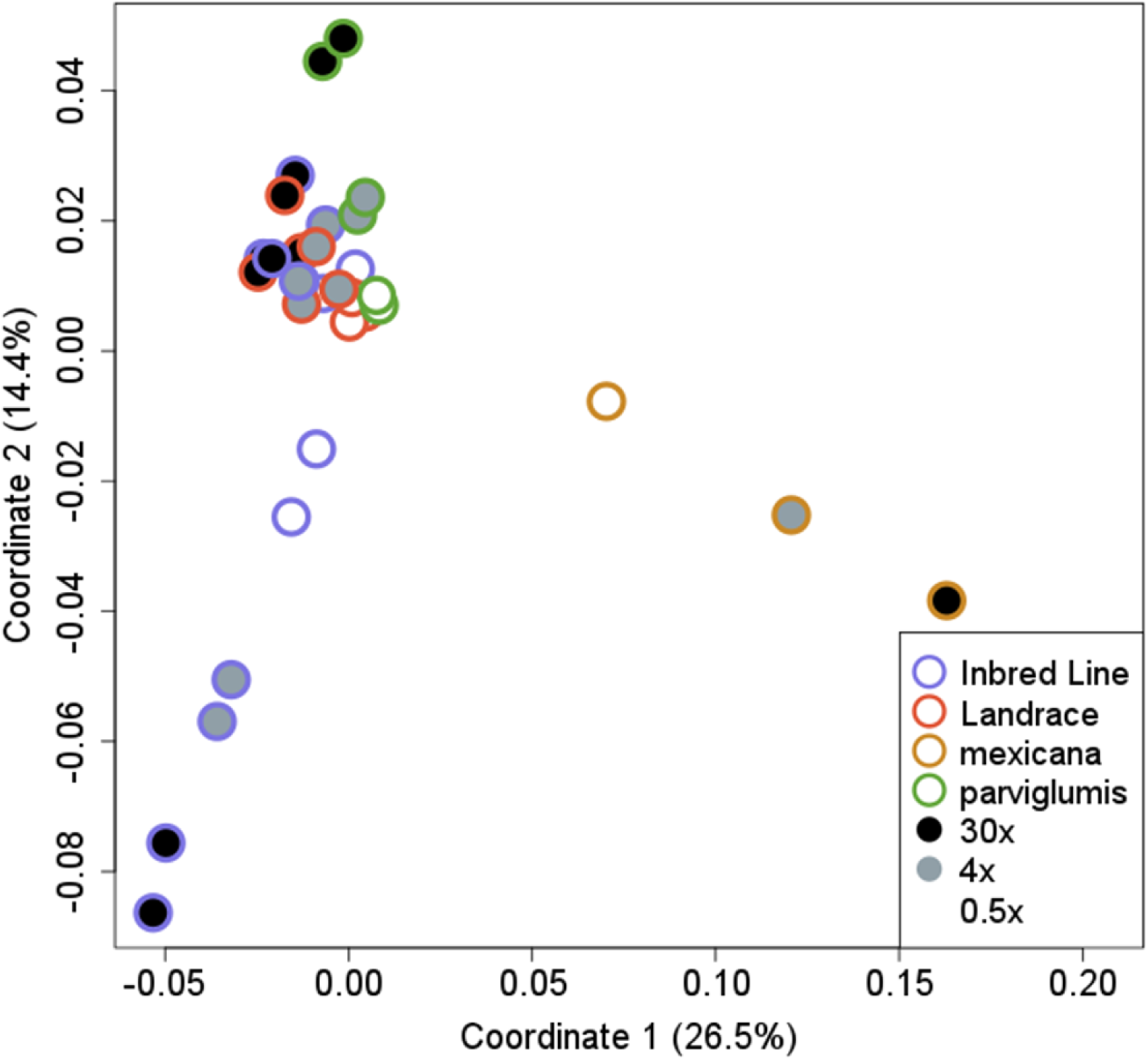
– Quantification of shared variance among downsampled maize accessions using the GATK variant caller. This figure presents a Multidimensional Scaling (MDS) plot featuring 11 downsampled individuals (*SI Appendix*, Table S2) at various sequencing coverages. The inner ring visually represents the average genome-wide sequencing coverage, with 30X coverage depicted in black, 3X coverage in gray, and 0.5X coverage in white. The outer ring provides information on the identity of the samples, including inbred lines (IL), outbred landraces (LR), as well as the maize progenitor subspecies *mexicana* and *parviglumis*.

**Figure S5.**
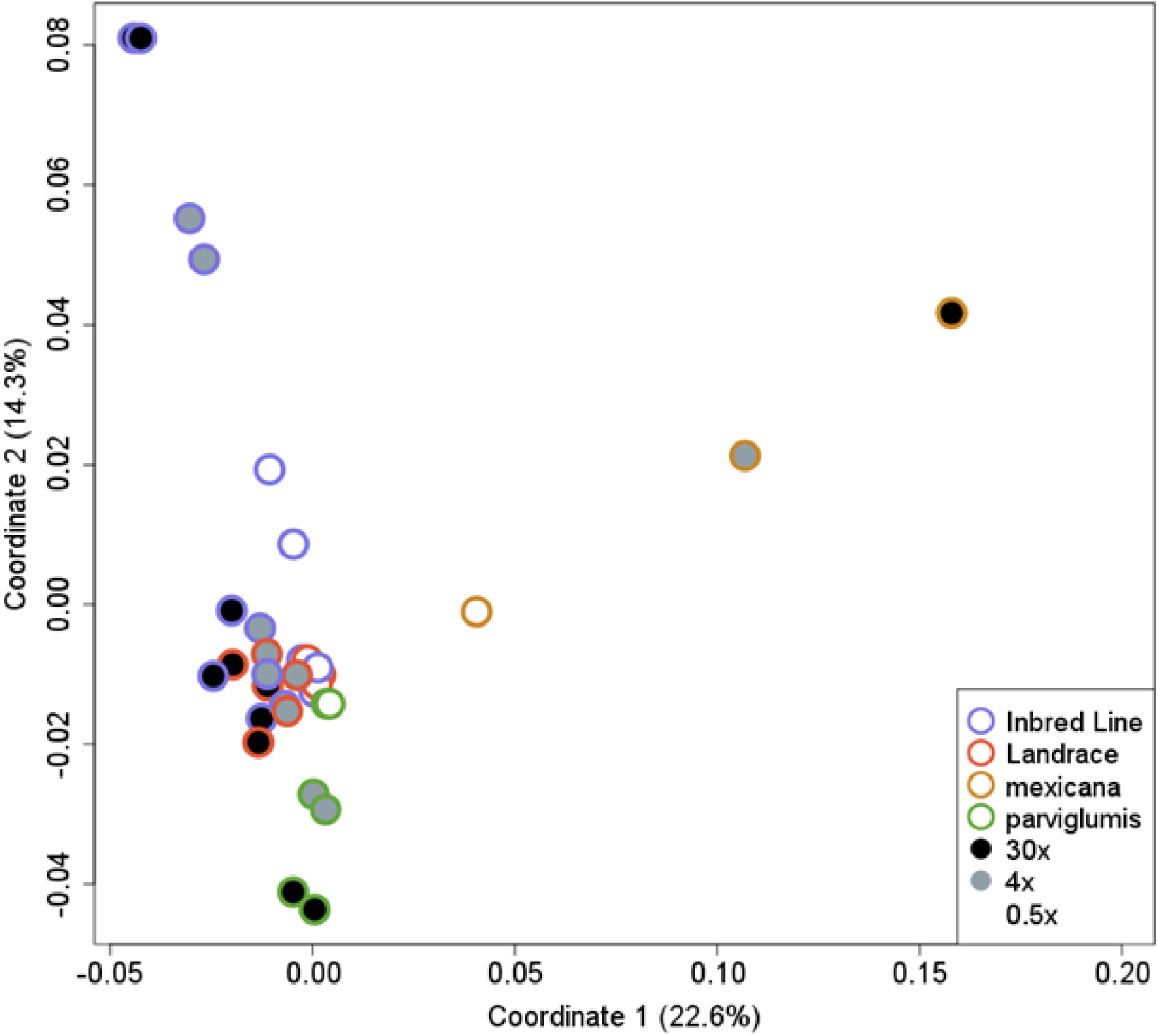
– Quantification of shared variance among downsampled maize accessions using the Samtools variant caller. This figure presents a Multidimensional Scaling (MDS) plot featuring 11 downsampled individuals (*SI Appendix*, Table S2) at various sequencing coverages. The inner ring visually represents the average genome-wide sequencing coverage, with 30X coverage depicted in black, 3X coverage in gray, and 0.5X coverage in white. The outer ring provides information on the identity of the samples, including inbred lines (IL), outbred landraces (LR), as well as the maize progenitor subspecies *mexicana* and *parviglumis*

**Figure S6.**
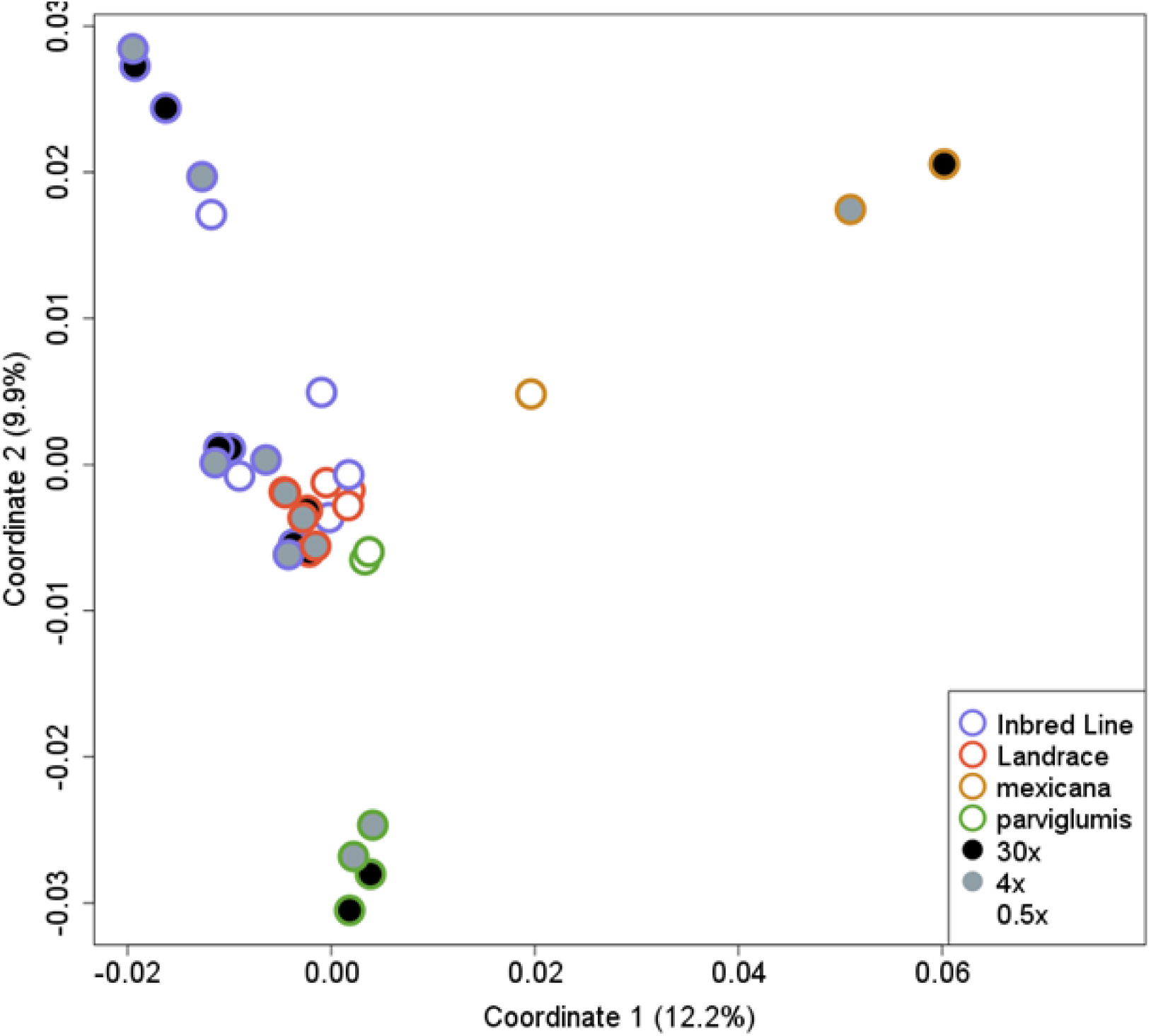
– Quantification of shared variance among downsampled maize accessions using the Freebayes variant caller. This figure presents a Multidimensional Scaling (MDS) plot featuring 11 downsampled individuals (*SI Appendix*, Table S2) at various sequencing coverages. The inner ring visually represents the average genome-wide sequencing coverage, with 30X coverage depicted in black, 3X coverage in gray, and 0.5X coverage in white. The outer ring provides information on the identity of the samples, including inbred lines (IL), outbred landraces (LR), as well as the maize progenitor subspecies *mexicana* and *parviglumis*.

**Figure S7.**
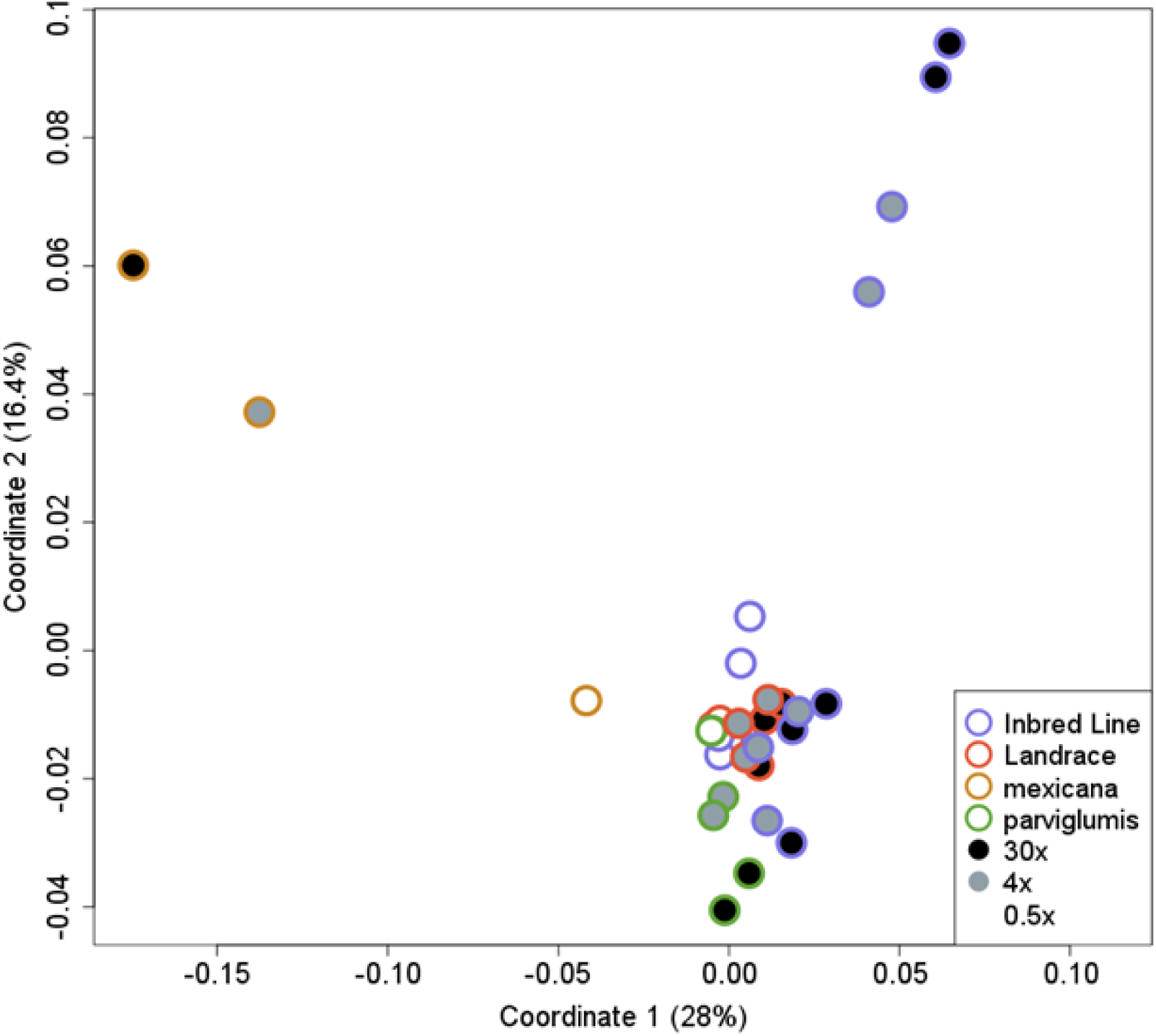
– Quantification of shared variance among downsampled maize accessions using the DeepVar variant caller. This figure presents a Multidimensional Scaling (MDS) plot featuring 11 downsampled individuals (*SI Appendix*, Table S2) at various sequencing coverages. The inner ring visually represents the average genome-wide sequencing coverage, with 30X coverage depicted in black, 3X coverage in gray, and 0.5X coverage in white. The outer ring provides information on the identity of the samples, including inbred lines (IL), outbred landraces (LR), as well as the maize progenitor subspecies *mexicana* and *parviglumis*

**Figure S8.**
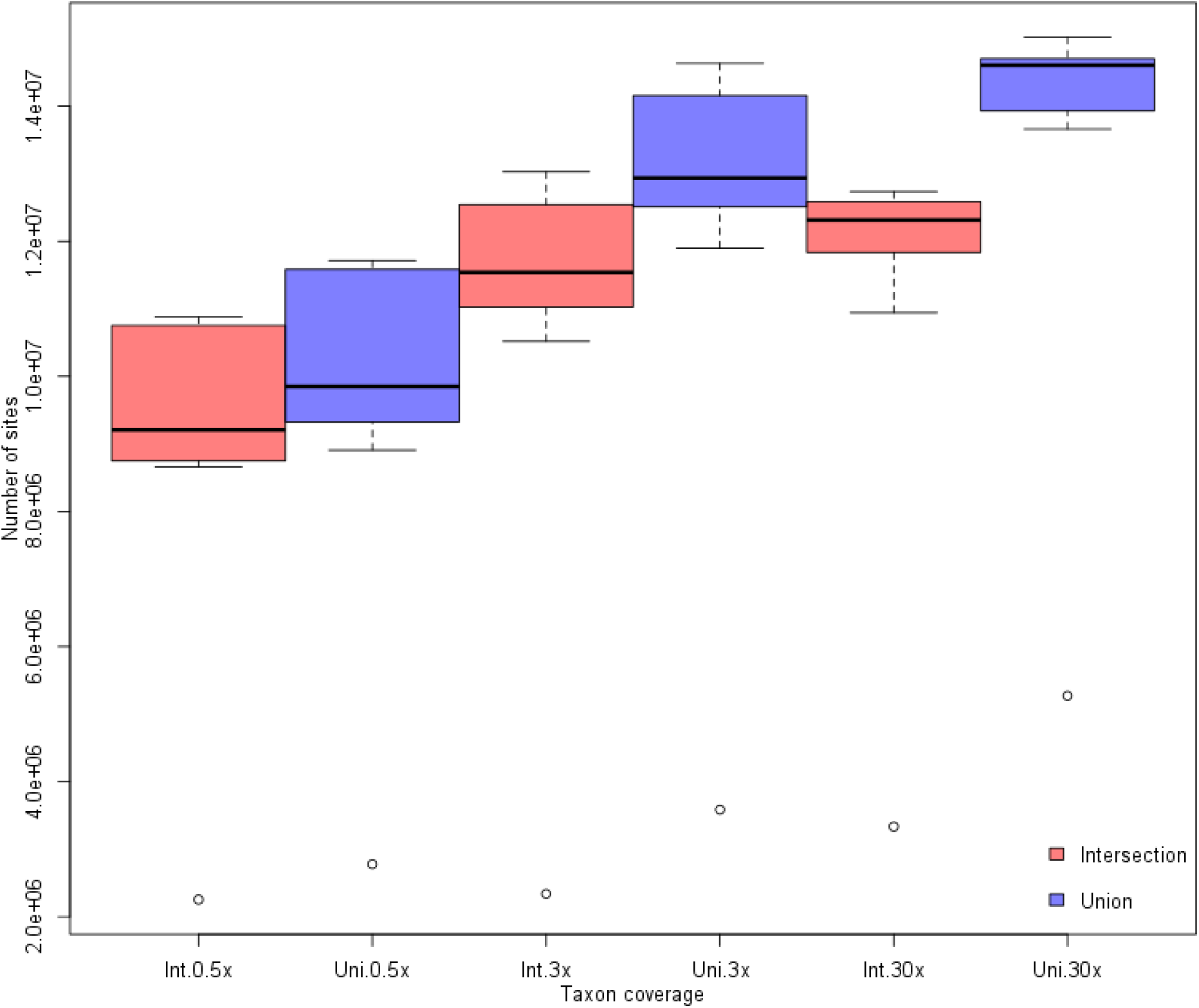
– Quantification of variant sites obtained through set operations at various sequencing coverages. This figure depicts boxplot distributions illustrating the outcomes of set operations, specifically the union (Uni) and intersection (Int), on variant sites. The Y-axis signifies the total count of identified variants with a Phred quality score greater than 0 for downsampled set individuals (*SI Appendix*, Table S2), while the X-axis represents different combinations of Intersection and Union strategies across sequencing depths of 0.5X, 3X, and 30X.

**Figure S9.**
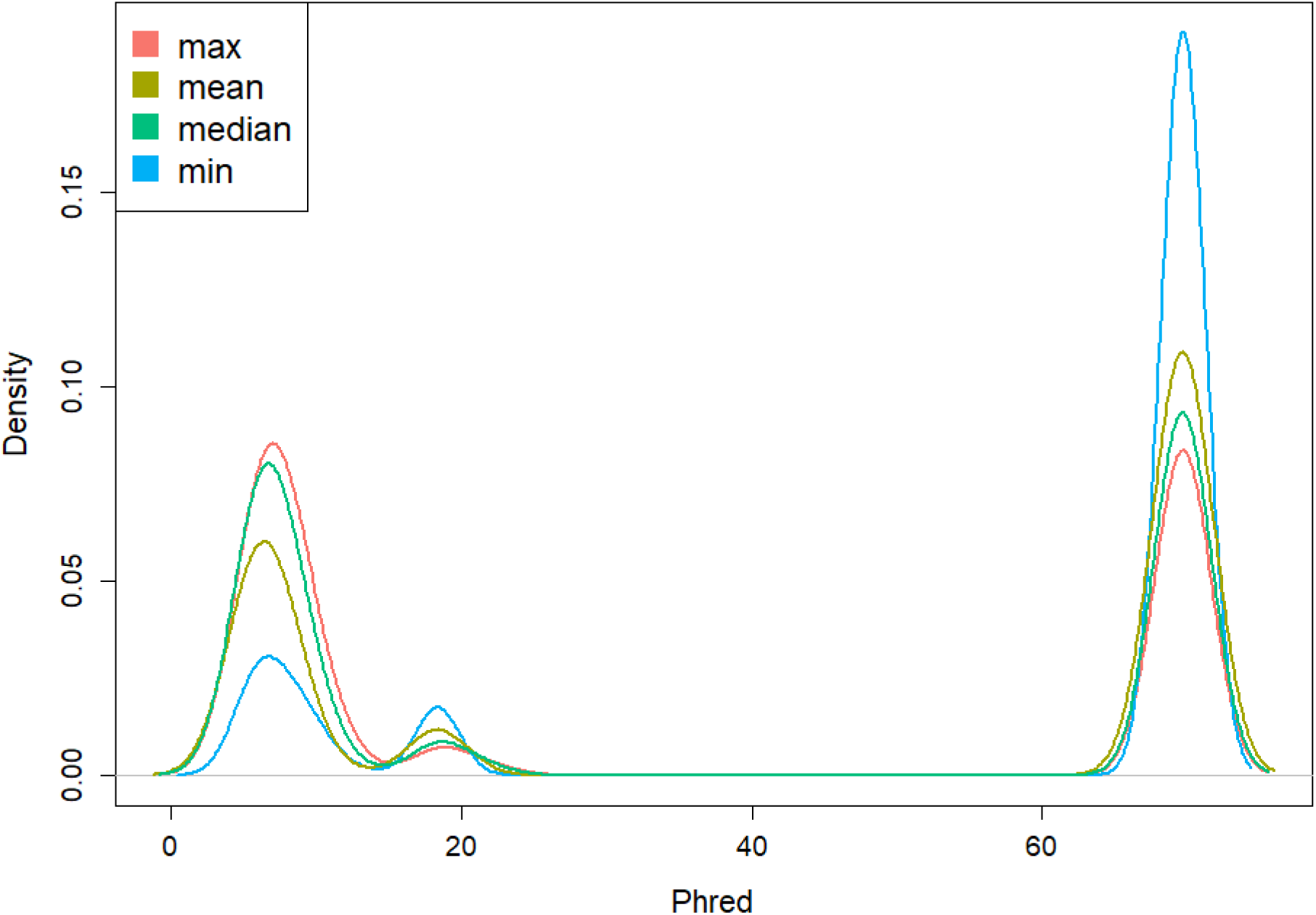
– Quantification of distribution profiles for Phred score values in variants identified by multiple callers, utilizing summary statistics. The X-axis presents the density function of Phred score profiles for various summary statistics, including maximum error (max), mean error (mean), median error (median), and minimum error (min) of the genotypes. These statistics are computed for maize accessions (SI Appendix, Table S3) at the same genomic position across different callers, thereby revealing the variability and characteristics of Phred scores within this context.

**Figure S10.**
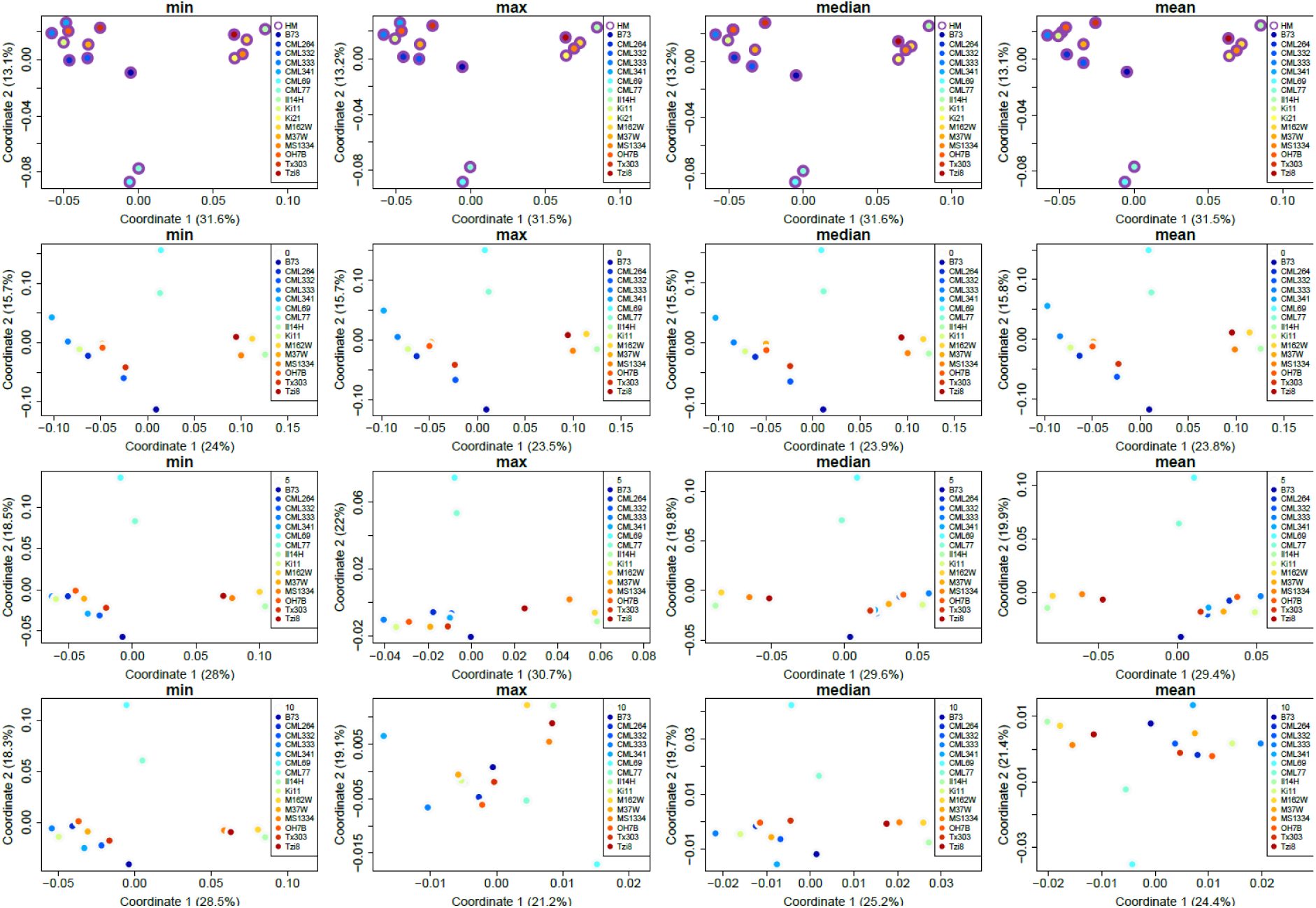
– Quantification of the impact of cutoffs on shared variance among maize accessions with variants identified by the UME caller compared to control. This figure showcases a series of Multidimensional Scaling (MDS) plots, each column presenting summary statistics computed for the same individual at identical genomic positions but detected by different variant callers and summarized into a single value. The summary statistics considered encompass maximum error (max), mean error (mean), median error (median), and minimum error (min) of genotypes. These MDS plots illuminate the variability and characteristics of error summary statistics in the context of variance among maize inbred line individuals. In the rows, information is organized regarding the quality filters applied during variant calling, starting with the control variants originating from HapMap3 in the first row. Subsequent rows detail the effects of cutoff values, including no cutoff (cutoff >= 0) in the second row, a cutoff of >= 5 in the third row, and a cutoff of >= 10 in the fourth row. The outer ring color-codes the control variants from HapMap3 in purple, contrasting with the white representation of the treatment effects under various combinatorial conditions. Additionally, the inner ring visually identifies each maize accession utilized in the analysis ( SI Appendix, Table S3).

**Figure S11.**
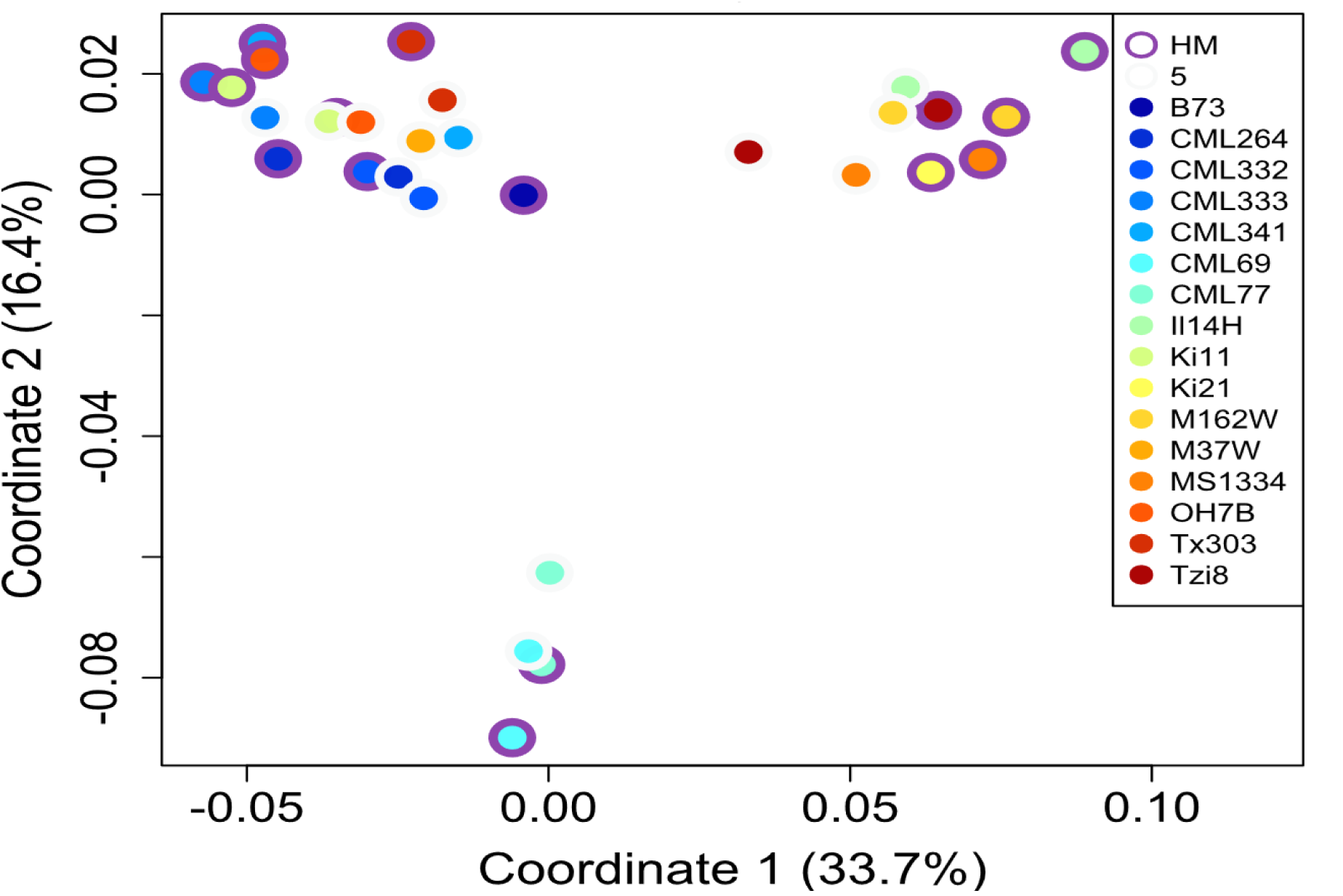
– Quantification of the impact of cutoff 5 on shared variance among maize accessions with variants identified by the UME caller compared to control. This figure presents a Multidimensional Scaling (MDS) plot, providing insights into the genetic variance among a series of maize accessions (*SI Appendix*, Table S3). The inner ring color-codes each individual maize accession for visual distinction. The outer ring differentiates control variants from HapMap3, depicted in purple, and the treatment effects of variant diversity obtained through the implementation of the UME calling strategy with a cutoff of 5. This treatment leverages the maximum error (max) summarization of Phred score variant probabilities to reveal the impact of the UME strategy on shared genetic variance within the maize accessions.

**Figure S12.**
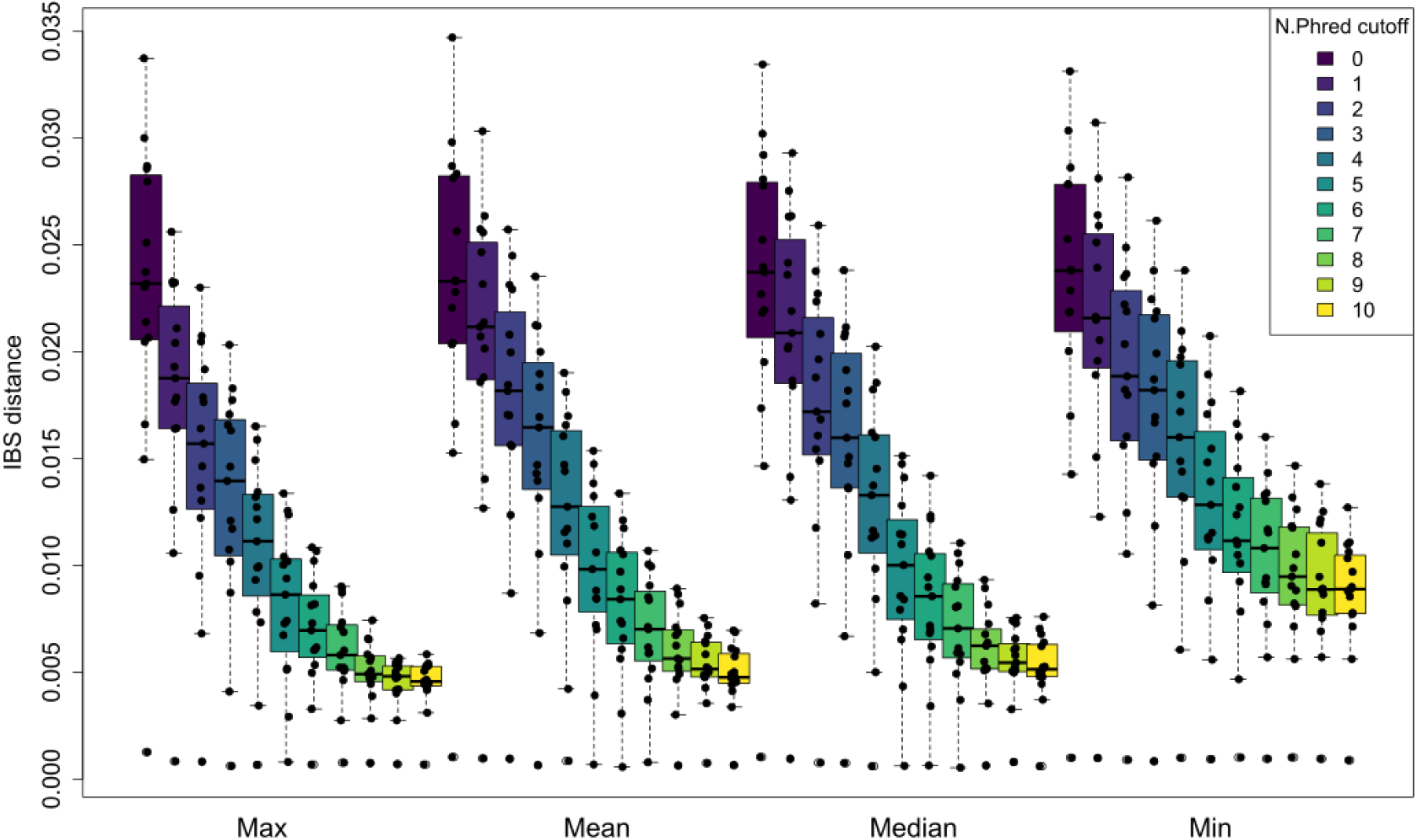
– Quantification of genetic distance between control variants and UME-called variants to assess the impact of summary statistics and variant error probability cutoffs. This figure presents box plots representing the distribution of Identity by State distance (IBS) among individual maize accessions (*SI Appendix*, Table S3). The analysis compares the control variant set from HapMap3 with its equivalent accession variant set called using the UME strategy under various combinations of summary statistics for modeling errors. These summary statistics include maximum error (max), mean error (mean), median error (median), and minimum error (min). Furthermore, different cutoff values are considered in this assessment. By examining the IBS distances, this figure provides insights into the genetic divergence to the control resulting from the choice of summary statistic and the application of different cutoff values in the context of variant error probabilities for maize accessions.

**Figure S13.**
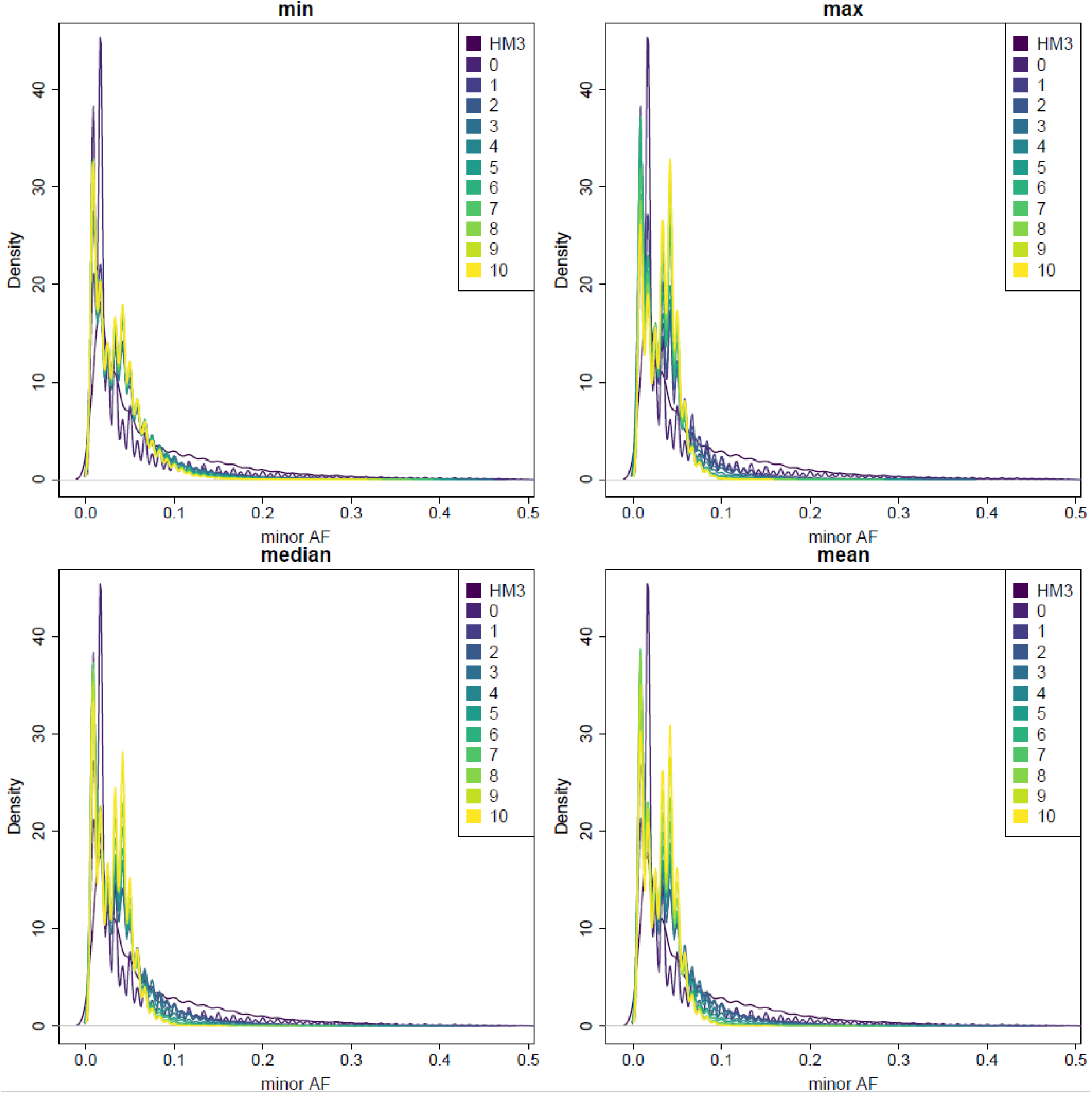
– Comparison of the impact of cutoffs and error summary statistics on the site frequency spectrum of UME-called variants. This figure features Folded Site Frequency Spectrum plots for variants obtained from maize accessions (*SI Appendix*, Table S3) using the UME variant caller. Control variants obtained from HapMap3 are represented in purple. Each panel within the figure corresponds to a distinct error summary strategy, including maximum error (max), mean error (mean), median error (median), and minimum error (min). The various colors within the panels indicate different error cutoff values. The X-axis illustrates the Minimum Allele Frequency (MAF), while the Y-axis represents the relative density of data per MAF value. These plots allow for a visual assessment of how the choice of error summary statistic and the application of different error cutoffs influence the site frequency spectrum of maize variants called by UME.

**Figure S14.**
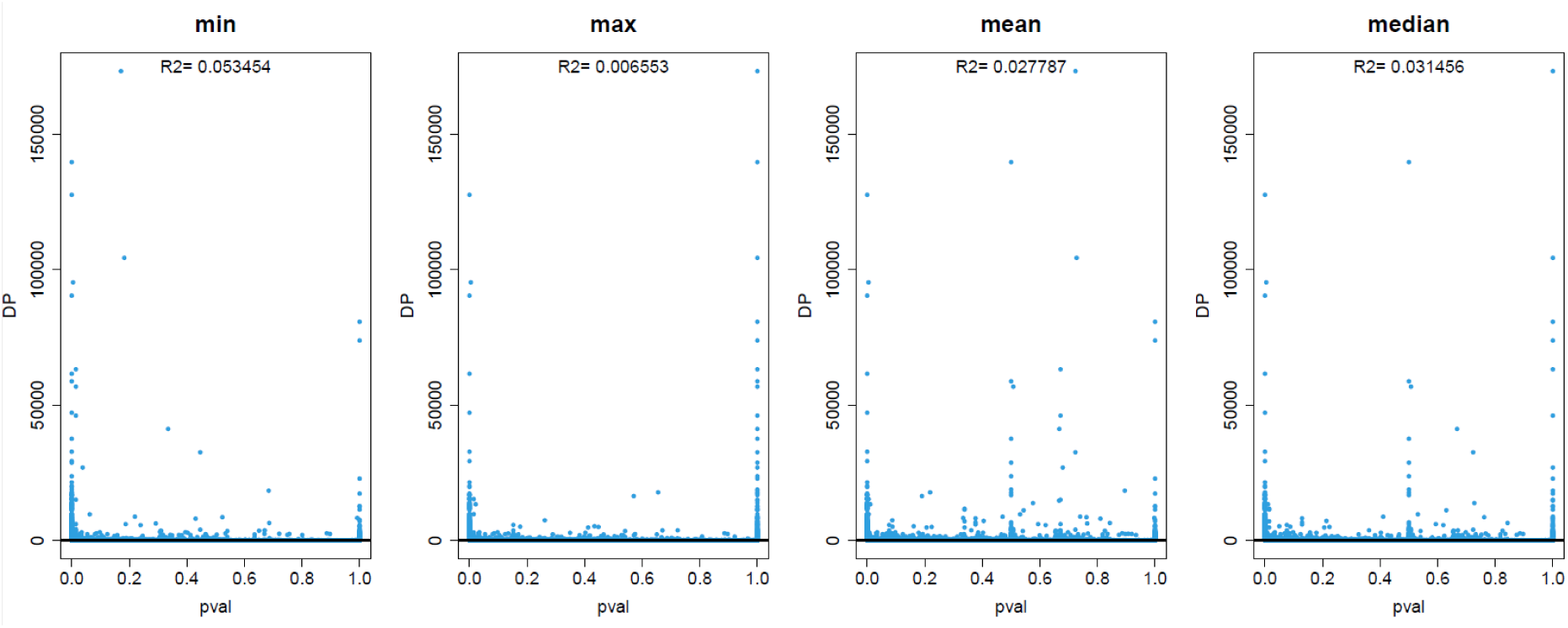
– Assessment of the relationship between sequencing depth and all P-values obtained by UME calling strategy with contrasting error summary statistics. This figure showcases scatterplots representing random samples of variants derived from maize accessions (SI Appendix, Table S3) obtained by UME caller, under different error summary statistics, including maximum error (max), mean error (mean), median error (median), and minimum error (min).In each scatterplot, the x-axis represents the sequencing depth, while the y-axis displays the associated p-values. The coefficient of determination (R^2) is reported to assess the extent to which p-values can be explained by sequencing depth.

**Figure S15.**
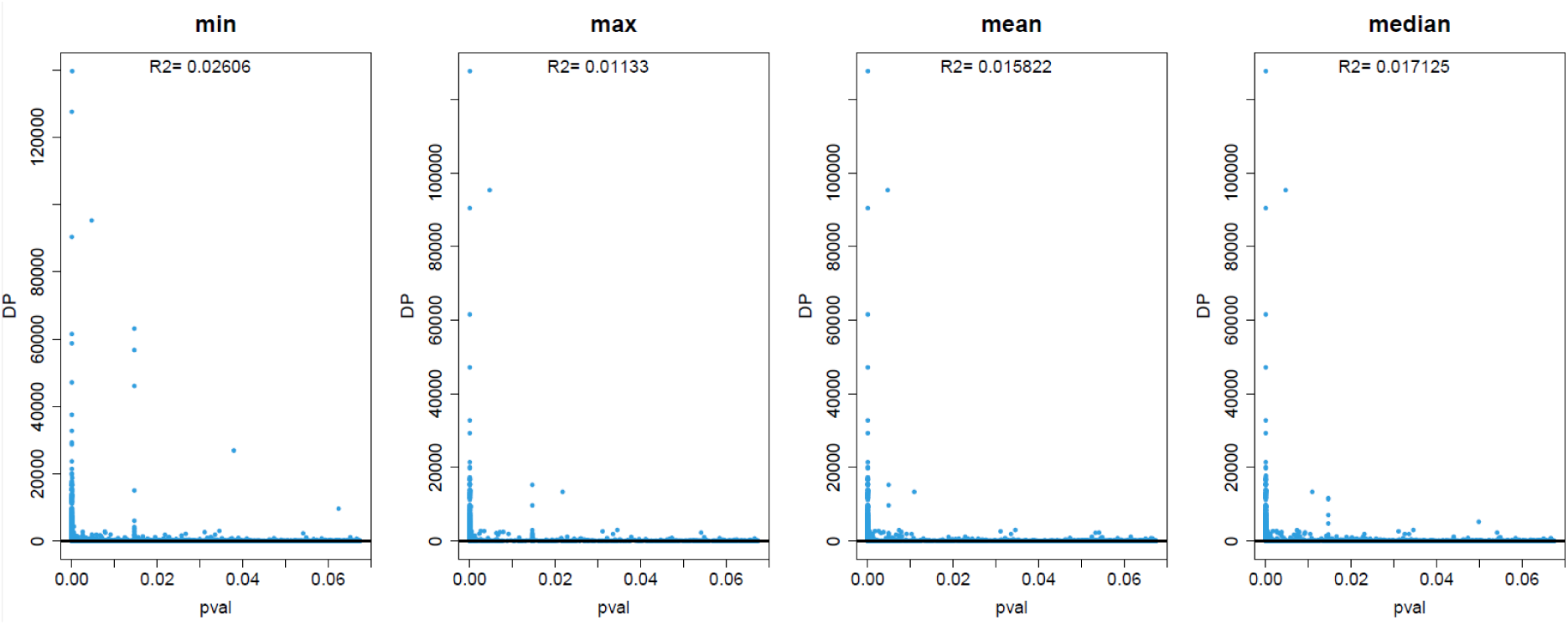
– Assessment of the relationship between sequencing depth and P-values obtained by UME calling strategy after a cutoff of 5 contrasting error summary statistics. This figure showcases scatterplots representing random samples of variants derived from maize accessions (SI Appendix, Table S3) obtained by UME caller, after a cutoff of 5, under different error summary statistics, including maximum error (max), mean error (mean), median error (median), and minimum error (min). In each scatterplot, the x-axis represents the sequencing depth, while the y-axis displays the associated p-values. The coefficient of determination (R^2) is reported to assess the extent to which p-values can be explained by sequencing depth.

**Figure S16.**
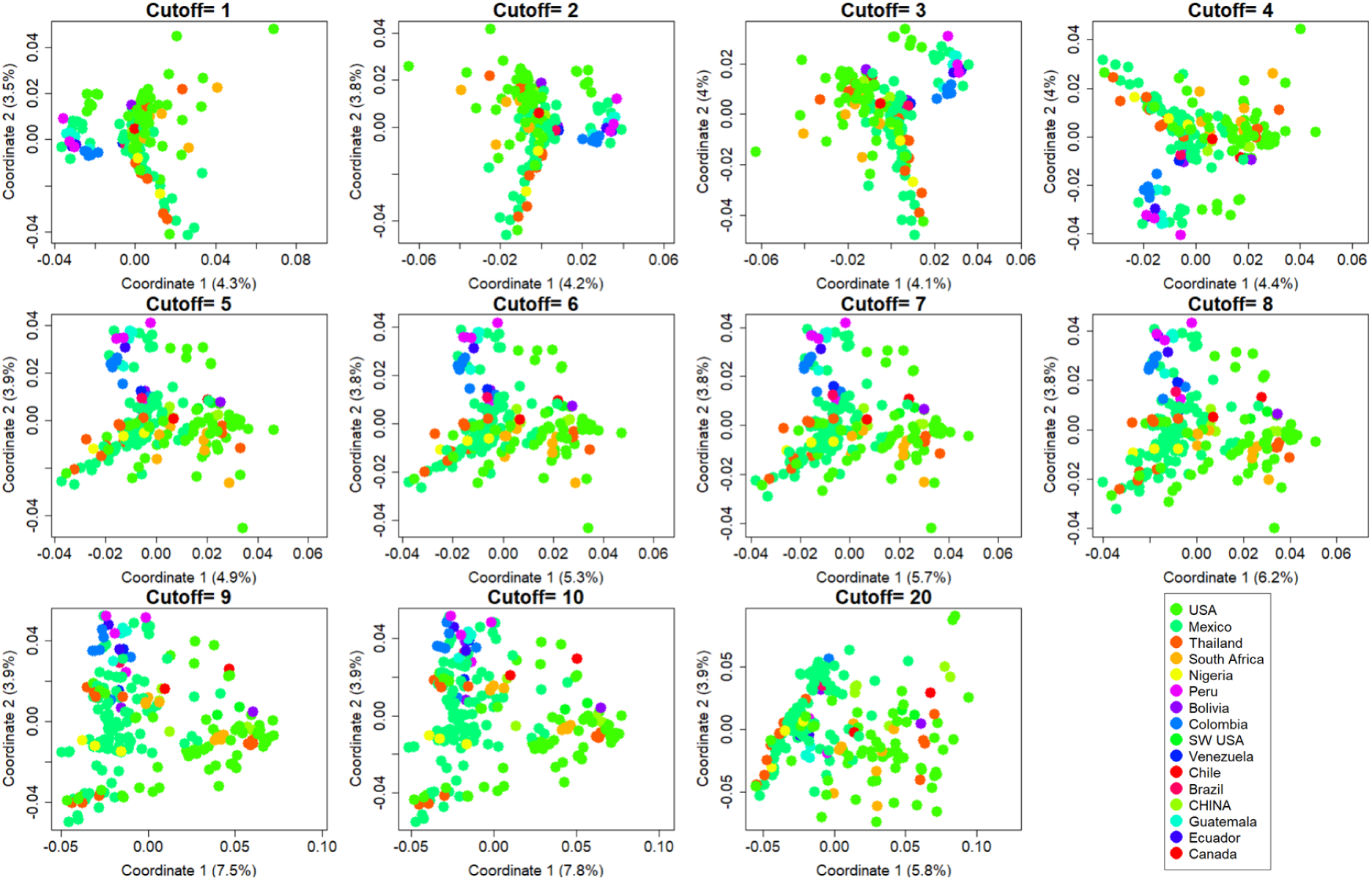
Assessment of the effect of error cutoff on variance explained by the coverage bias-controlled haplotype map database of balanced maize diversity, called by UME. This figure presents a series of Multidimensional Scaling (MDS) plots, with each panel representing the variation explained by variants obtained using the UME variant caller. The analysis employs a maximum error summary statistic while systematically varying the error cutoff applied to the resulting variants. The maize accessions used for this analysis are sampled from a specified dataset (SI Appendix, Table S1). Each individual in the MDS plots is color-coded according to their respective country of origin, as indicated in the legend provided in the last panel.

**Figure S17.**
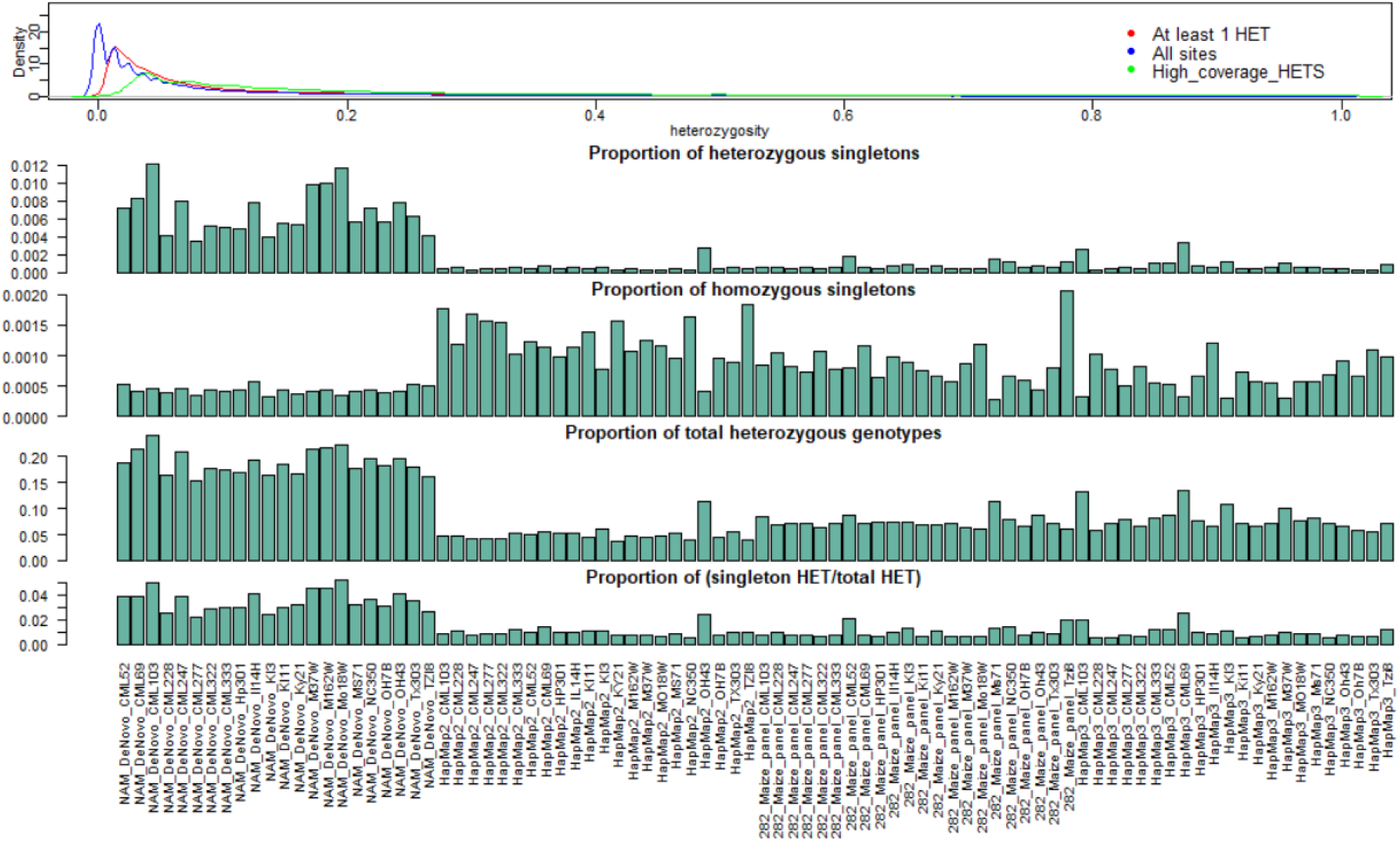
– Heterozygosity profiling of inbred lines within HapMap variant call using UME.

**Figure S18.**
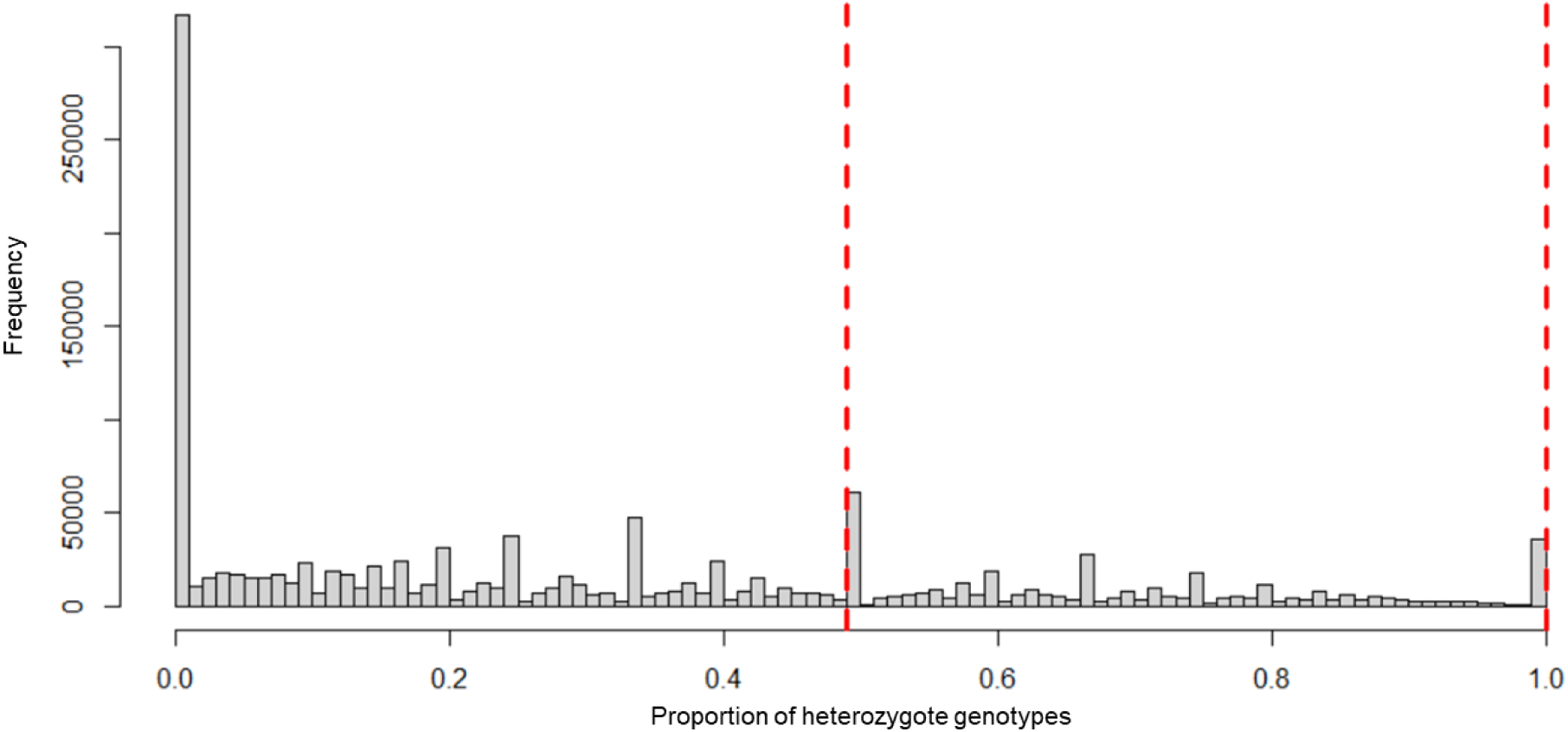
– Distribution of the proportion of unbalanced allelic depths in heterozygous calls over sites. Red lines represent the range of high proportion unbalanced heterozygous sites marked as paralogs.

**Figure S19.**
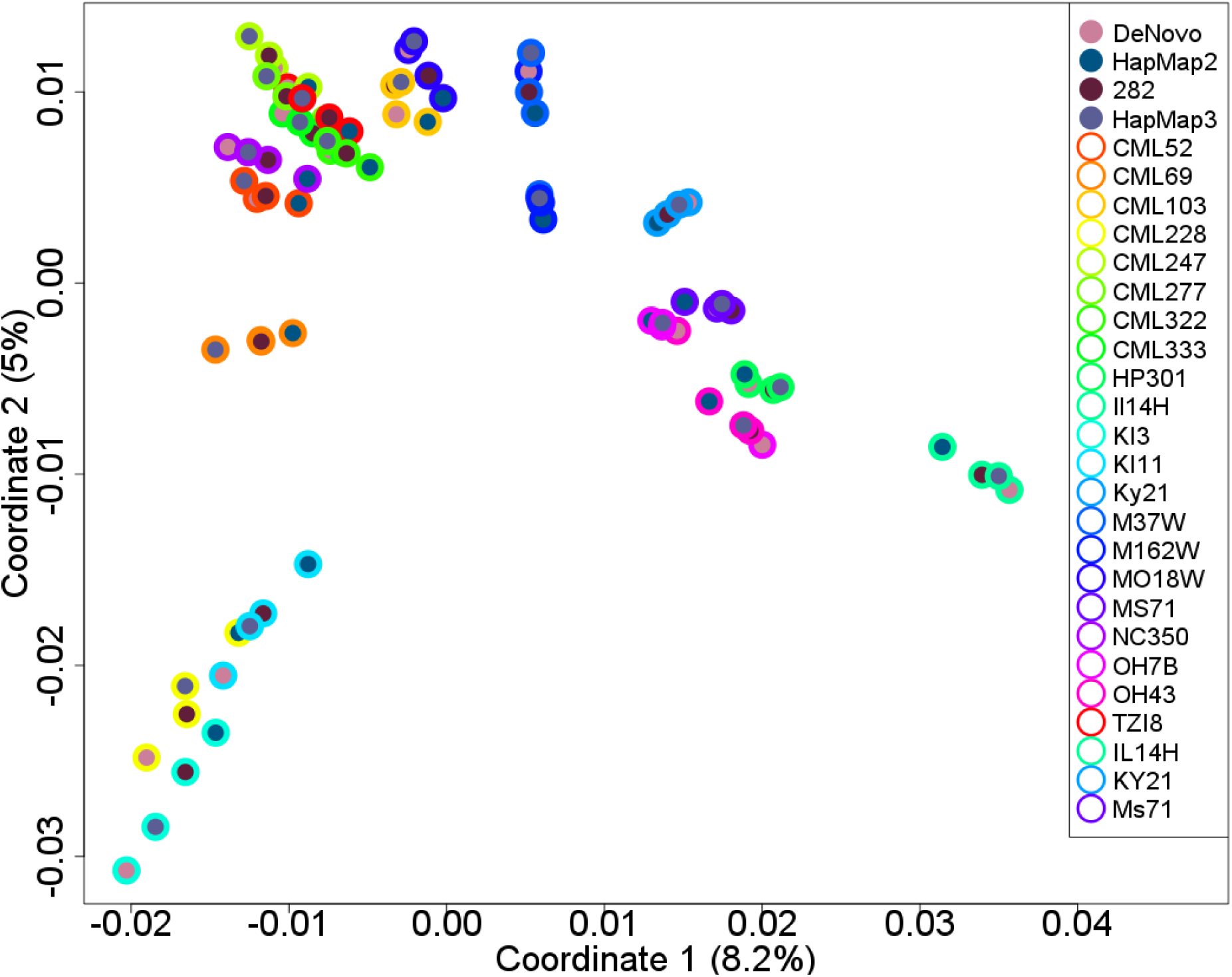
– Quantification of consistency in shared variance obtained by UME among individuals from the same maize accession across different sequencing efforts. This figure presents a Multidimensional Scaling (MDS) plot, showcasing maize inbred lines that are common across four distinct sequencing efforts (*SI Appendix*, Table S1): NAM founders de novo assembly (DeNovo), maize genetic association panel 282 (282), HapMap2, and HapMap3. All variants from these four sequencing efforts were identified using the UME variant caller, employing a maximum error summary statistic.

## Notes

### Competing Interest Statement

The authors have declared no competing interest.

